# *Stk11* deficiency promotes tumor CXCL16 secretion and a CXCL16-dependent expansion of *Areg^pos^ CXCR6^pos^* Vγ6 γδT17 in lung cancer

**DOI:** 10.64898/2026.11.25.769235

**Authors:** Katherine J. Horrigan, Joshua M. Ragusa, Somen K. Mistri, Oliver Dienz, Brianna M. Hilton, Sean M. Lenahan, Allison R. Racela, Megan Tarte, Paula B. Deming, Yvonne M.W. Janssen-Heininger, David J. Seward, Matthew E. Poynter, Jonathan E. Boyson

## Abstract

IL-17-producing gamma delta (γδT17) T cells can promote tumor growth and metastasis. How the tumor microenvironment controls the activation, expansion, and function of γδT17 in different cancers remains unclear. Using an autochthonous model of *Kras^G12D^/Stk11*-driven lung adenocarcinoma coupled with scRNAseq and V(D)J profiling, we found that *Kras^G12D^Stk11*-loss tumors induced a rapid expansion of lung CXCR6^+^ Vγ4 and amphiregulin-expressing CXCR6^+^ Vγ6 γδT17 cells. Single-cell immune profiling of the *Kras^G12D^Stk11*-loss tumors revealed a prominent *Krt8*^+^ alveolar epithelial cell (KAC) tumor subtype enriched in a number of immune-relevant genes, including *Cxcl16* and its sheddase *Adam10*, and *in vitro* culture revealed that KACs, but not their normal epithelial cell counterparts, secreted CXCL16. Finally, we found that *in vivo* blockade of CXCL16 inhibited the expansion of lung Vγ6^+^ γδT17 in response to *Kras^G12D^Stk11*-loss tumors. Together, these data suggest that *Stk11* loss promotes a CXCL16-secreting *Krt8^+^* KAC tumor subtype, and that CXCL16-CXCR6 interactions are necessary to drive the expansion of pro-tumor γδT17 cells in lung cancer.

## INTRODUCTION

γδ T cells are unconventional innate-like T cells that are enriched in mucosal tissues like the lung, gut, and skin, where they play critical roles in tumor immunity, infectious disease, and tissue homeostasis. A defining characteristic of γδ T cells is their ability to rapidly produce large amounts of cytokines, chemokines, and growth factors that allow them to shape both the magnitude and quality of the developing innate and adaptive immune responses. In the setting of cancer, γδ T cells have previously been demonstrated to exert both pro-tumor and anti-tumor functions (Mallu et al., 2026; Reis et al., 2022; Silva-Santos et al., 2019). This contrasting role has been attributed to their different functional subsets, with IFNγ-producing γδT1 cells being associated with anti-tumor immunity (Gao et al., 2003; Girardi et al., 2001; Lanca et al., 2013; Weifeng He, 2010), while IL-17-producing γδT17 cells are associated with tumor growth and metastasis (Coffelt et al., 2015; Jin et al., 2019; Li et al., 2014; Rei et al., 2014; Wu et al., 2014).

Lung cancer is the leading cause of cancer-related deaths worldwide, and the vast majority of these cancers consist of non-small cell lung cancer (NSCLC) (Bray et al., 2024; Zhang et al., 2023). Despite recent treatment advances, the overall survival rate for NSCLC remains low (Herbst et al., 2018). NSCLCs exhibit significant heterogeneity at both the cellular and genetic levels, contributing to substantial variability in the tumor-immune microenvironment (Biton et al., 2018; Chen et al., 2014; Skoulidis et al., 2015; Skoulidis et al., 2018). Cancers containing loss-of-function mutations in the *STK11* (*LKB1*) tumor suppressor, which frequently co-occur in *KRAS*-driven lung adenocarcinomas (Skoulidis et al., 2015), rapid tumor growth is associated with impaired PD-L1 expression (Ji et al., 2007; Skoulidis et al., 2015), increased tumor-associated neutrophils, and decreased αβ T cell infiltration (Koyama et al., 2016; Schabath et al., 2016; Skoulidis et al., 2015), resulting in relative resistance to immunotherapy (Skoulidis et al., 2018).

In the setting of lung cancer, γδ T cells have been found to be enriched in human lung tumors (Wu et al., 2022b), and in mouse models they have been demonstrated to exert significant effects on both the tumor microenvironment and on tumor burden (Jin et al., 2019; Raffo-Iraolagoitia et al., 2026). In an autochthonous *Kras^G12D^/p53^fl^* model of lung adenocarcinoma, commensal microbiota-driven IL-1β and IL-23 production drove lung Vγ4 and Vγ6 γδT17 cell proliferation, which resulted in increased tumor burden via the recruitment of immunosuppressive tumor-associated neutrophils to the lung (Jin et al., 2019). In a *Kras^G12D^/Myc^fl^* model of lung cancer, increased numbers of lung Vγ1 T cells induced the expansion of profibrotic airway macrophages (Raffo-Iraolagoitia et al., 2026). Whether the lung γδ T cell compartment is influenced by the genetic drivers of the tumor remains largely unexplored (Biton et al., 2018; Busch et al., 2016; Koyama et al., 2016; Schabath et al., 2016; Skoulidis et al., 2018), and the mechanisms that govern the expansion, survival, and persistence of γδ T cells within the tumor microenvironment remain unclear.

Here, we set out to explore these questions using a novel model of *Kras^G12D^/Stk11^fl^*-driven lung adenocarcinoma. We report that rapid disease progression in this model was associated with the expansion of the pro-tumoral Vγ4 and *Areg*^+^ *Cxcr6*^+^ Vγ6 γδT17 subsets in the lungs. Moreover, single-cell profiling of the *Kras^G12D^/Stk11^fl^*tumors revealed an expansion of *Krt8*^+^ alveolar intermediate cells that expressed both the CXCR6 ligand, *Cxcl16,* and the sheddase *Adam10*. Additionally, we found that *Stk11*-deficient tumor epithelial cells, but not their normal epithelial cell counterparts, secreted CXCL16. Finally, we found that *in vivo* blockade of CXCL16-CXCR6 interactions was sufficient to prevent the expansion of lung Vγ6 γδT17 in response to tumor. Together, these data identify CXCL16 as a critical mediator of the expansion of pro-tumoral γδT17 subsets in the *Kras^G12D^/Stk11^fl^* lung tumor-immune microenvironment.

## MATERIALS AND METHODS

### Mice

B6.Scgb1a1^Cre/+^.Rosa26^LSL-tdTomato/+^. Kras^LSL-G12D/+^.Stk11^+/+^ (K), B6.Scgb1a1^Cre/+^. Rosa26^LSL-tdTomato/+^.Kras^LSL-G12D/+^.Stk11^flox/flox^ (KS), B6.Scgb1a1^Cre/+^.Rosa26^LSL-tdTomato/+^.Kras^+/+^.Stk11^+/+^ (WT), and B6.Scgb1a1^Cre/+^.Rosa26^LSL-tdTomato/+^.Kras^+/+^.Stk11^flox/flox^ (STK11) mice (Snoke et al., 2025) were bred at the University of Vermont. Mice were given *ad libitum* access to chow and water and housed in a specific pathogen-free animal facility on a 14:10 light:dark cycle. Tumor formation was induced in 8–10-week-old animals by intra-peritoneal (i.p.) administration of 2 mg of tamoxifen (Sigma-Aldrich #T5648) in corn oil for 3 consecutive days. Male and female mice were used in approximately equal numbers in all experiments. For survival assays, mice administered tamoxifen were observed daily to monitor survival. All animal procedures were carried out with approval from the University of Vermont Institutional Animal Care and Use Committee.

For CXCL16 blocking experiments, tumor induction was initiated with 2 mg of tamoxifen in corn oil on days 0, 1, and 2 as described above. Mice were treated with 30 μg intranasal (i.n.) and 150 μg i.p. of anti-CXCL16 antibody (BioXCell #BE0450) three times a week for two weeks starting on day 5 and were euthanized on day 20 for analysis.

### Tissue histochemistry and pathology

Lung tissue was fixed by placing harvested lung lobes in tissue cassettes and incubating overnight at 4°C in PBS/4% PFA. The tissue was then placed in 70% ethanol for long-term storage. Tissues were dehydrated, embedded in paraffin, and 8 μM sections were stained with hematoxylin and eosin. Tumor burden was assessed by a board-certified pathologist (D.J.S.) as previously described (Snoke et al., 2025).

### Tissue processing for flow cytometry

Single-cell suspensions were prepared from lungs as described (Benoit et al., 2015). Briefly, lungs were minced with scissors in digestion buffer containing DMEM, 1 mg/ml collagenase IV (Gibco), and 0.2 mg/ml DNase (Sigma-Aldrich). Lung tissue was incubated in a shaking incubator at 200 RPM at 37°C for 20 minutes, triturated with a 16-gauge blunt needle, followed by an additional 20-minute incubation at 200 RPM at 37°C. Red blood cells were lysed using Gey’s solution. Finally, the digested lung tissue was filtered through 70-μm mesh and washed with PBS/2% FBS. Cells were counted on either a MACSQuant10 following propidium iodide staining or with a hemacytometer after trypan blue staining. Single-cell suspensions from spleen and lymph node were obtained by passing tissue through 70-μm nylon mesh. Splenic red blood cells were lysed using Gey’s solution.

### *In vitro* cytokine stimulation

Lung leukocytes were enriched using the EasySep^TM^ Mouse CD45 Positive Selection Kit (STEMCELL #18945) according to the manufacturer’s instructions. CD45^+^ lung cells were then incubated with 25 ng/ml PMA (Sigma P8139) and 500 ng/ml ionomycin (Sigma I-0634) in Click’s media/5% FBS/1X penicillin/1X streptomycin/4 mM L-Glutamine/50 μM 2-ME for 2 hours at 37°C, 5% CO_2_, before adding 2 μM monensin (Sigma M5273) and incubating for an additional 3 hours.

### Flow cytometry

Single-cell suspensions were first incubated with LIVE/DEAD^TM^ Fixable Blue Dead Cell stain (Thermo Fisher Scientific) for 30 minutes at 4°C in PBS. Cells were then washed and resuspended in Fc Block (BioLegend) in PBS/2% FBS/0.1% sodium azide. After a 10-minute incubation at 4°C, fluorophore-conjugated antibodies specific for cell-surface markers were added and incubated for an additional 30 minutes at 4°C. After washing, cells were fixed in PBS/1% PFA for at least 30 minutes and then stored in PBS/2% FBS/0.1% sodium azide at 4°C, prior to data collection.

For nuclear staining, cells underwent staining for cell surface markers as described above, followed by fixation and permeabilization with Foxp3 transcription factor fixation/permeabilization buffer (ThermoFisher Scientific) for 30 minutes at 4°C, after which they were washed and incubated with 25 μg/ml rat IgG prior to adding antibodies against transcription factors. For intracellular cytokine staining, cells were fixed with PBS/1% PFA, permeabilized with PBS/1% FCS/0.1% sodium azide/0.1% saponin, then incubated with 25 μg/ml rat IgG before adding cytokine-specific antibodies.

Flow cytometry data was collected on a Cytek® Aurora (Cytek Biosciences) 4 laser (355/405/488/640) spectral cytometer, and data analysis was performed using FlowJo (BD Biosciences). Antibodies used in these experiments were: anti-BrdU-APC (BU20A), anti-CD4-BUV395 (RM4-5), anti-CD8α-BUV563 (53-7.3), anti-CD11b-Alexa488 (M1/70), anti-CD11b-PE (M1/70), anti-CD11b-APC/Cy7 (M1/70), anti-CD11c-Alexa488 (N418), anti-CD11c-PE (N418), anti-CD19-Alexa488 (6D5), anti-CD19-APC/Cy7 (1D3), anti-CD25-Alexa488 (PC61), anti-CD44-BV480 (IM7), anti-CD44-APC/Cy7 (IM7), anti-CD45-eFluor506 (30-F11), anti-CD45-BUV395 (30-F11), anti-CD45.2-Alexa488 (104), anti-CD45RB-BUV496 (C363-16A), anti-CD45RB-APC/Cy7 (C363-16A), anti-CD64-APC (X54-5/7.1), anti-CD103-BV605 (2E7), anti-CD103-PE/Fire700 (QA17A24), anti-CD170-BV480 (1RNM44N), anti-CD170-Alexa647 (S17007L), anti-CD186-SparkRed718 (SA051D1), anti-CD198-BV711 (SA214G2), anti-CD279-BV711 (29F.1A12), anti-CD279-PE/Cy7 (29F.1A12), anti-CD326-APC (G8.8), anti-Foxp3-Alexa647 (FJK-16s), anti-IFNγ-BV650 (XMG1.2), anti-IFNγ-Alexa647 (XMG1.2), anti-IL17A-BV605 (TC11-18H10.1), anti-ST2-BV421 (DIH9), anti-KLRG1-BV605 (2F1/KLRG1), anti-Ly6C-eFluor450 (HK1.4), anti-Ly6C-PerCP/Cy5.5 (HK1.4), anti-Ly6G-PacificBlue (1A8), anti-Ly6G-PerCP/Cy5.5 (1A8), anti-MHCII-Alexa700 (M5/114.15.2), anti-NK1.1-PE/Cy7 (PK136), anti-Rorγt-BV650 (Q31-378), anti-Tbet-PE/Dazzle594 (4B10), anti-TCRβ-BV785 (H57-597), anti-TCRγδ-BV421 (GL3), anti-TCRγδ-Alexa488 (GL3), anti-TCRγδ-PE (GL3), anti-TCRγδ-PE/CF594 (GL3), anti-TCRγδ-Alexa647 (GL3), anti-TCR-Vγ1-BV421 (2.11), anti-TCR-Vγ1-APC (2.11), anti-TCR-Vγ4-PE/Cy7 (UC3-10A6), anti-TCR-Vγ5-FITC (536), anti-TCR-Vγ6-PE (1C10-1F7), anti-Vγ6-RB705 (1C10-1F7), anti-TCR-Vδ4-PE (GL2), and anti-TCR-Vδ6.3-R718 (8F4H7B7).

### *In vivo* cell proliferation assay

Mice were injected i.p. with 1 mg BrdU (Sigma #B5002) for 3 consecutive days. 1-2 hours after the final BrdU injection, mice were euthanized, and lungs were harvested for flow cytometric analysis, as described above. Incorporation of BrdU in γδ T cells was assessed according to the manufacturer’s instructions (BrdU Flow Kit; BD Biosciences).

### Lung cytokine/chemokine profiling

Lungs were harvested and homogenized in PBS/0.5% IGEPAL with protease inhibitor cocktail (Thermo Scientific PIA32953). Homogenized tissue samples were then centrifuged at 4°C at 1500 x g for 15 minutes, and the supernatant was collected and stored at -80°C. Total protein concentration was calculated using the Pierce BCA kit (Thermo Fisher Scientific A55864). Protein detection was conducted using multiplexed Discovery Assays (Luminex® 200™ platform, Eve Technologies). Protein concentrations were normalized based on total lung protein content.

### Cell sorting and library preparation for gd T cell CITE-seq and V(D)J enrichment

For γδ T cell scRNAseq analysis, single-cell suspensions were prepared from lungs 28 days post-tamoxifen induction using collagenase/DNAse as described above. Single-cell suspensions were stained with anti-TCRδ-PE antibody and enriched using anti-PE microbeads kit (Miltenyi Biotec) according to the manufacturer’s instructions. Following viability staining with Fixable Viability Dye eFluor^TM^ 780 (Thermo Fisher Scientific), cells were surface stained with fluorescent antibodies against CD45.2, CD11b, CD11c, CD19, and TCRβ, in addition to oligonucleotide-conjugated hashtag antibodies (TotalSeqC-BioLegend). TCRδ^+^ cells were sorted from the CD11b^-^ CD11c^-^ CD19^-^ TCRβ^-^ population of CD45.2^+^ live cells on a BD FACS Aria 3. Live γδ T cells from B6.Scgb1a1^Cre/+^.Rosa26^LSL-tdTomato/+^.Kras^LSL-G12D/+^.Stk11^+/+^ and B6.Scgb1a1^Cre/+^.Rosa26^LSL-tdTomato/+^.Kras^LSL-G12D/+^.Stk11^flox/flox^ lungs or B6.Scgb1a1^Cre/+^.Rosa26^LSL-tdTomato/+^.Kras^+/+^.Stk11^+/+^ and B6.Scgb1a1^Cre/+^.Rosa26^LSL-tdTomato/+^.Kras^+/+^.Stk11^flox/flox^ lungs (n = 4 mice per strain) were pooled and captured (about 2000 cells/mouse) using a 10X Chromium controller (10X Genomics), according to the manufacturer’s instructions.

For lung epithelial cell scRNAseq analysis, single-cell suspensions from lungs on day 23 post-tamoxifen induction were prepared as described above, except that the enzymatic digestion mix consisted of DMEM, 0.5 mg/ml collagenase IV, 0.5 mg/ml elastase, and 0.2 mg/ml DNase. Digested lung tissue was filtered through 70-μm mesh, washed with PBS/2% FBS, and CD45^+^ cells were depleted from each sample using the EasySep^TM^ Mouse CD45 Positive Selection Kit (Stemcell Technologies #18945) according to the manufacturer’s instructions. Cells were then stained with Fixable Viability Dye eFluor^TM^ 780 (Thermo Fisher Scientific) followed by fluorescent antibodies against CD45.2 and EpCAM, and oligonucleotide-conjugated hashtag antibodies (TotalSeqC, BioLegend). Live EpCAM^+^, CD45.2^-^, tdTomato^+^ cells were sorted on a BD FACS Aria 3. Equal numbers of cells from B6.Scgb1a1^Cre/+^.Rosa26^LSL-tdTomato/+^.Kras^LSL-G12D/+^.Stk11^+/+^ and B6.Scgb1a1^Cre/+^.Rosa26^LSL-tdTomato/+^.Kras^LSL-G12D/+^.Stk11^flox/flox^ lungs or B6.Scgb1a1^Cre/+^. Rosa26^LSL-tdTomato/+^. Kras. ^+/+^Stk11^+/+^ and B6.Scgb1a1^Cre/+^.Rosa26^LSL-tdTomato/+^.Kras^+/+^.Stk11^flox/flox^ lungs (n = 2 mice per strain) were pooled and captured (about 8000 cells) using a 10X Chromium controller.

### Library preparation for tumor single-cell RNA sequencing

Library amplification and preparation for γδ T cells and lung epithelial cells was conducted using a NextGEM Single Cell 5’ Reagent Kit v2 (Dual Index) according to the manufacturer’s instructions (10X Genomics). For γδ T cells, a V(D)J enrichment library was also prepared, with modifications as previously described (Mistri et al., 2024). After library quality and concentration were checked using a BioAnalyzer, libraries were pooled, and sequenced (paired-end 26 x 91 bp) on a Singular Genomics G4 sequencer.

### CITE-seq quality control, filtering, and demultiplexing

Analysis of single-cell CITE-seq datasets was conducted as previously described (Mistri et al., 2024). Briefly, raw sequence reads were processed using CellRanger software v6.1.0 (10X Genomics) and mapped to the GRCm38 (mm10) mouse reference genome. For the γδ T cell data sets, clustering analysis was conducted only after removing all TCR transcripts using the “cellranger reanalyze” argument of Cell Ranger software. Quality control, filtering, UMAP projections, and clustering analyses were conducted using Seurat (v5.2.0) R package (Hao et al., 2021). Briefly, cells with a low number of unique genes (<500 for the γδ dataset and the tumor dataset), a frequency of mitochondrial reads >5%, and a frequency of ribosomal protein genes <5% were eliminated from further analysis. In some analyses, cell cycle genes were regressed out during the data normalization step. For the γδ dataset, one cluster of cells expressing macrophage-specific genes was manually removed.

For each dataset, scTransform was used to normalize the gene expression data (Hafemeister and Satija, 2019). After principal component analysis, we performed unsupervised clustering of cells using specific numbers of top PCs (γδ = 50, tumor = 30) and resolution values (γδ = 1.2, tumor = 0.8) to produce a Uniform Manifold Approximation and Projection (UMAP) for each dataset. The gene expression data was then visualized using the scCustomize R package (Marsh, 2021). The differentially expressed genes for each cluster were identified using FindAllMarkers() function using the Wilcoxon Rank Sum test. To identify differentially expressed genes between mouse strains, we performed a pseudobulk differential expression (DE) analysis of aggregated read counts using the DESeq2 package (Love et al., 2014), which accounts for biological replicates (Squair et al., 2021).

To analyze the V(D)J library sequences, Cell Ranger v7.1.0 (‘cellranger vdj’ argument) was used to map the TCR V(D)J reads to a custom-made reference containing sequences from the IMGT database (Lefranc, 2011). The Cell Ranger V(D)J output was then analyzed with a custom R script. Briefly, the TRGV and TRAV/TRDV contigs identified by Cell Ranger V(D)J were demultiplexed to individual barcodes, after which the dominant TRGV and TRAV/DV transcripts were selected per cell. Finally, the V(D)J information of each cell was integrated with scCITE-seq data and visualized using a combination of Seurat and custom-made R scripts (Mistri et al., 2024). We note that the methods used here cannot adequately assess the true TRGV2 repertoire. Therefore, we report it here for accuracy but have omitted TRGV2 from our analysis.

### Lung epithelial cell culture and chemokine analysis

Primary cell culture of lung epithelial cells was conducted by preparing single-cell suspensions from lungs day 28 post-tamoxifen induction as described above using digestion buffer containing DMEM, 0.5 mg/ml collagenase IV, 0.5 mg/ml elastase, and 0.2 mg/ml DNase. Digested lung tissue was filtered through 70-μm mesh, washed with PBS/2% FBS, after which CD45+ cells were depleted from each sample using the EasySep^TM^ Mouse CD45 Positive Selection Kit (Stemcell Technologies #18945) according to manufacturer’s instructions. CD45-depeleted cells were then stained with Fixable Viability Dye eFluor^TM^ 780 (Thermo Fisher Scientific) followed by fluorescent antibodies against CD45.2 and CD326 (EpCAM). Live EpCAM^+^ CD45.2^-^, tdTomato^+^ cells were sorted on a BD S8 spectral sorter. Cells were washed, resuspended in DMEM with 5% FBS, 100 units pen/strep, 4 mM L-glutamine, 0.25 μg/ml Fungizone, 0.1 μg/ml cholera toxin, 10 μg/ml insulin, 5 μg/ml transferrin, 30 μg/ml BPE, 25 ng/ml EGF, and 0.01 μM retinoic acid, and then incubated at 37°C 5% CO_2_ on plates that had been previously coated overnight with 100 mg/ml of collagen in 0.02N glacial acetic acid.

Cell culture supernatants were collected after 5 days of culture and analyzed for CXCL16 protein levels via Luminex® (R&D systems # LXSAMSM) according to manufacturer’s instructions.

### Differential gene expression in human lung epithelial cells

RNAseq FASTQ files from our previous study in which *STK11* was deleted from H2009 human lung epithelial cells (Lenahan et al., 2024) were aligned using Salmon v2.5.1 (Patro et al., 2017), and differential gene analysis was performed using DESeq2 (Love et al., 2014), as previously described (Mistri et al., 2024).

### Statistical Analysis

All statistical analyses were conducted using Prism (GraphPad Software). Unpaired Student t-tests, one-way ANOVA with Dunnett’s T3 test, or two-way ANOVA with Sidak multiple comparisons corrections were used where appropriate. The Kaplan-Meier method with log-rank (Mantel-Cox) test was used for survival studies. Statistical tests were considered significant when p-values were less than 0.05. Error bars represent standard deviation unless otherwise indicated.

## RESULTS

### Tumor-intrinsic Stk11 loss is associated with an expansion of IL-17-producing lung Vγ4 and Vγ6 T cells

To investigate the function and regulation of γδ T cells in lung adenocarcinoma (LUAD), we employed a recently developed autochthonous mouse model of *Kras^G12D^*-driven lung adenocarcinoma with a concurrent deletion of the tumor suppressor *Stk11* (Snoke et al., 2025). In this model, *Scgb1a1*-driven Cre recombinase drives expression of the *Kras^G12D^* allele with or without the concurrent deletion of *Stk11* (referred to as KS and KRAS strains, respectively) primarily in club cells. The inclusion of an LSL-tdTomato allele in the *Rosa* locus allows the identification and fate-mapping of Cre-expressing epithelial cells. A comparison of disease progression between KRAS and KS mice revealed that while expression of the *Kras^G12D^* allele resulted in increased mortality compared to wild-type controls, the concurrent deletion of *Stk11* in conjunction with *Kras^G12D^* expression resulted in further increased mortality (Fig. 1A) and increased tumor burden (Fig. 1B) over *Kras^G12D^* alone, consistent with previous studies (Koyama et al., 2016; Nagaraj et al., 2017). The rapid disease progression observed upon *Stk11* loss in *Kras^G12D^* mice was associated with significant alterations in the lung tumor microenvironment, with notable increases in CD4 αβ T cells, Foxp3^+^ Tregs, iNKT cells, and an increased DC2/DC1 ratio (Fig. S1). Consistent with a previous report (Koyama et al., 2016), we observed a significant increase in neutrophils in KS lungs, including a population of Ly6G^+^SiglecF^+^ neutrophils that have been associated with a pro-tumor microenvironment in lung adenocarcinoma (Pfirschke et al., 2020), and we observed significant increases in both IL-6 and G-CSF in KS lungs (Fig. S1).

**Figure 1.**
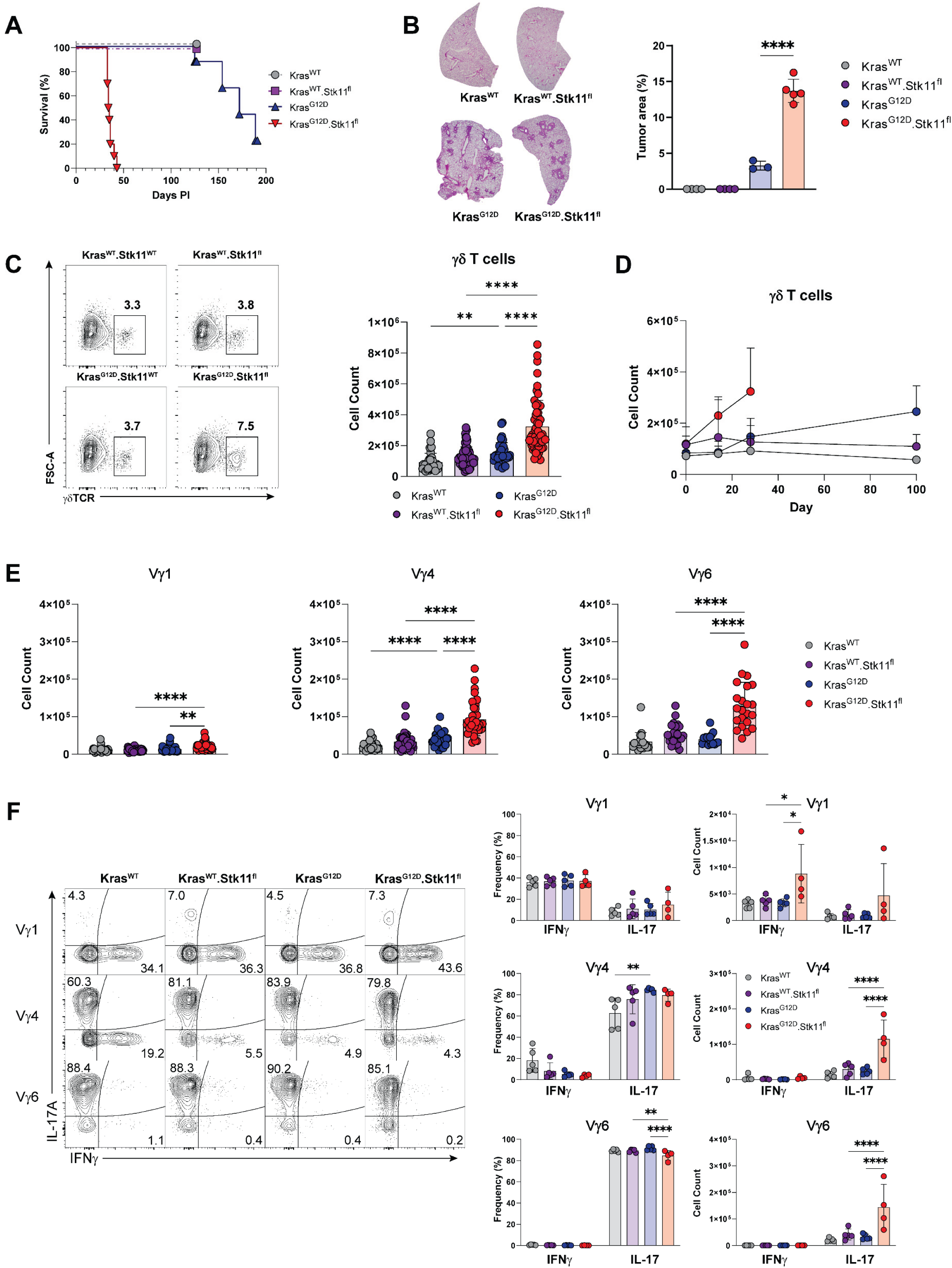
Tumor-intrinsic loss of *Stk11* is associated with increased mortality, increased tumor burden, and rapid expansion of lung γδT17 cells. **A.** Survival analysis of *Kras^WT^* (n = 8), *Kras^WT^.Stk11^fl^*(n = 8), *Kras^G12D^* (n = 8) *Kras^G12D^.Stk11^fl^*(n = 10) mice after tamoxifen induction. Kaplan-Meier method with log-rank (Mantel-Cox) test, p < 0.0001. **B.** Tumor burden analysis among the four genotypes 30 days after tamoxifen induction. Representative H&E sections are shown at (*left)*, while cumulative tumor area data are shown at (*right)*. **C.** Representative contour plots of lung γδ T cells (*lef*t), and cumulative lung γδ T cell counts at day 28 post tamoxifen induction (*right)* D. Time course of lung γδ T cell counts from *Kras^WT^*, *Kras^WT^.Stk11^fl^*, *Kras^G12D^,* and *Kras^G12D^.Stk11^fl^* mice between days 0 – 100 after tamoxifen induction. **E.** Numbers of individual lung γδ T cell subsets 28 days post tamoxifen induction. **F.** Preferential expansion of lung γδT17 after tumor *Stk11* loss. 28 days after tamoxifen induction, lung leukocytes were stimulated *ex vivo* with PMA and ionomycin, then stained for intracellular cytokine production. Representative contour plots (*left*) and cumulative data (*right)* are shown. One-way ANOVA with Dunnett’s T3 multiple comparisons test. *p < 0.05, **p < 0.01, ***p < 0.001, **** p < 0.0001.

In addition to the changes noted above, a significant increase in γδ T cell number was present in the lungs of KS mice as compared to their KRAS and control counterparts (Fig. 1C), with significant increases noted as early as 14 days after induction of tumorigenesis (Fig. 1D). Further examination revealed that Vγ4 and Vγ6 subsets accounted for the vast majority of these increases and that the number of lung Vγ1 T cells, while also significantly increased in KS mice, accounted for only ∼10% of the lung γδ

T cells (Fig. 1E). Examination of lung γδ T cell cytokine production revealed a significant expansion of IL-17-producing Vγ4 and Vγ6 T cells in the lungs of KS mice such that these γδT17 cells accounted for the vast majority of the increased number of γδ T cells in KS lungs (Fig. 1F). We also noted that there was a significant increase in the number of IFN-γ-producing Vγ1 T cells in the KS lung, although they comprised only a fraction of the total lung γδ T cell population, Together, these data suggest that club cell-intrinsic *Stk11* loss induces rapid disease progression and significant changes in the lung tumor-immune microenvironment that includes a significant expansion of lung γδT17 T cells.

### Single-cell V(D)J profiling reveals the expansion of an *Areg*+ population of Vγ6 γδT17 cells in KS lungs

To investigate the functional diversity of the lung γδ T cells expanding as a result of tumor *Stk11* loss, we next conducted single-cell RNA sequencing paired with V(D)J profiling on γδ T cells sorted from the lungs of all four strains (n=4 each strain; Fig. S2). Because KS mice begin to succumb to disease 32 – 40 days after tumor induction, we chose to sort γδ T cells 28 days after tumor induction. Unsupervised clustering revealed the presence of 3 “superclusters” of lung γδ T cells--a γδT17 supercluster enriched in the canonical markers *Il17a, Il17f,* and *Rorc,* consisting of 9 unique clusters, a γδT1 supercluster enriched in *Ifng*, *Tbx21, Klrd1, Klrc1*, and *Klrc2* consisting of 3 unique clusters, and a naïve γδ T cell cluster enriched for *Cd24a, Ccr7, and Sell* (Figs. 2A-2C, Suppl. Table 1).

**Figure 2.**
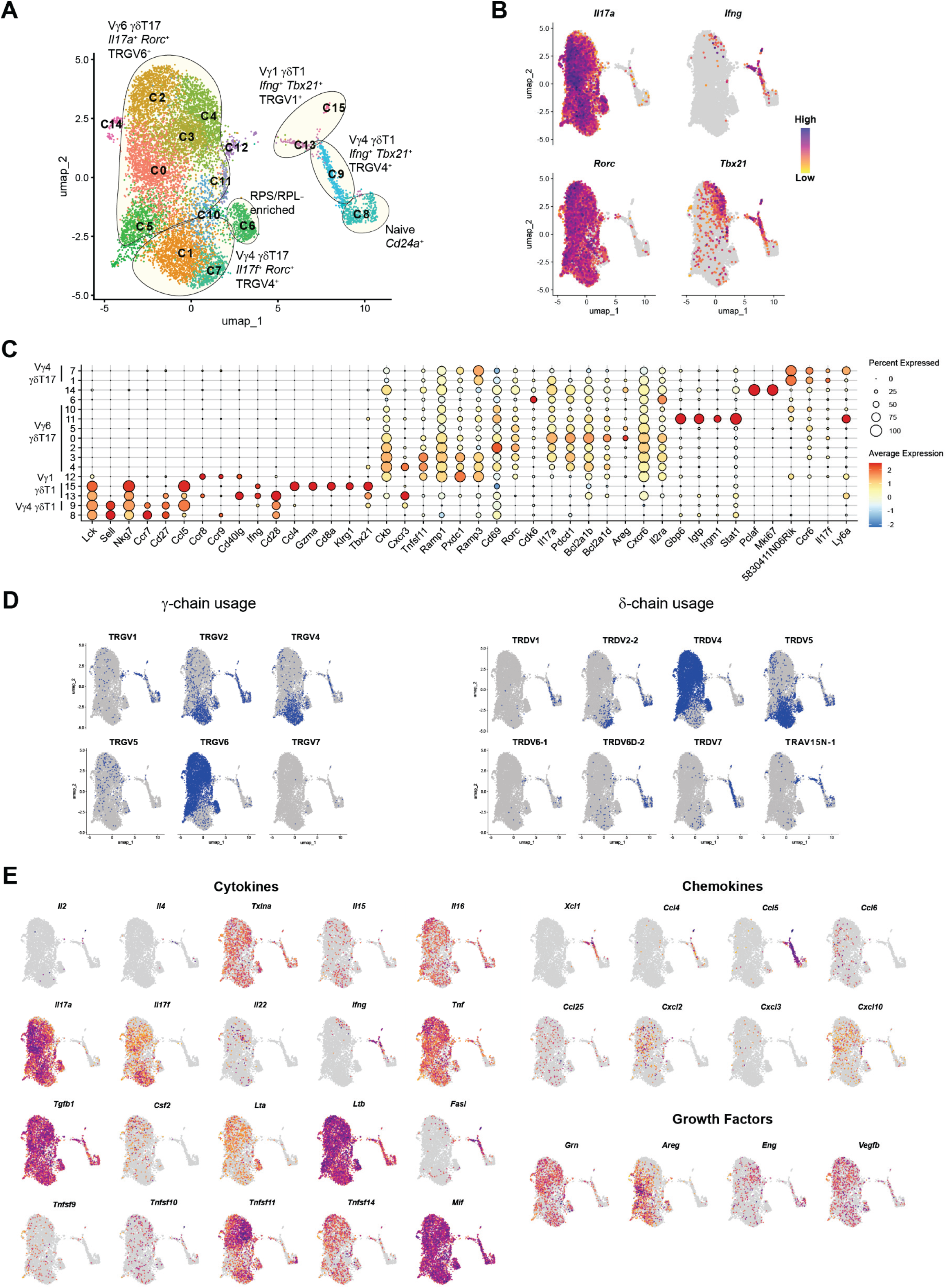
scCITE-seq analysis of lung γδ T cells reveals high Vγ6 transcriptional diversity. **A.** UMAP visualization of sorted lung γδ T cells pooled from *Kras^WT^* (WT), *Kras^WT^.Stk11^fl^* (STK11), *Kras^G12D^* (KRAS), and *Kras^G12D^.Stk11^fl^* (KS) (n = 4 each genotype) mice 28 days after tamoxifen induction. **B.** Select feature plots depicting canonical γδT1 (*Tbx21, Ifng*) and gdT17 *(Il17a, Rorc*) markers. Each cell is color-coded based on gene expression (purple: high, yellow: low). **C.** Dot plot illustrating the scaled expression levels of selected cluster-defining genes. Gene expression levels are normalized and depicted using a color scale ranging from low (blue) to high (red). The size of the dot represents the percentage of cells in the indicated cluster expressing the gene. **D.** UMAP representations of lung γδ T cells showing the TRGV (*left*) and TRDV (*right*) chain V-segment usage in blue. **E.** Feature plots of selected cytokine, chemokine, and growth factor genes (purple: high, yellow: low).

V(D)J profiling revealed that the γδT17 clusters were divided into 7 TRGV6 clusters (C0, C2, C3, C4, C5, C10, and C11) and 2 TRGV4 clusters (C1 and C7). Examination of the TRGV4 clusters revealed that γδ T cells with paired TRGV4 and TRDV5 chains (Vγ4/Vδ5) were present in both the C1 and C7 clusters, but that γδ T cells with paired TRGV4 and TRDV2-2 chains (Vγ4/Vδ4) were largely restricted to the C7 cluster (Fig. 2D). Examination of the TRDV CDR3 sequences paired with TRGV4 sequences revealed a prominent germline-encoded TRDV5 sequence previously associated with γδT17 (Kashani et al., 2015; Mistri et al., 2024) (Fig. S2). Further investigation of the TRGV4 γδT17 clusters revealed that C1 was enriched for *Itgb1*, *Il17a*, *Pdcd1*, and *Ctla4*, while C7 was enriched for *Ly6a, Socs3,* and *Dusp10* (Fig. 2C). Consistent with these observations, we found that Vγ4/Vδ4^-^, but not Vγ4/Vδ4^+^, cells exhibited increased CTLA4 and PD-1 expression in KS lungs (Fig. S2), suggesting that Vγ4/Vδ5 but not Vγ4/Vδ4 T cells may be preferentially activated in KS lungs. In addition, we noted small but significant populations of TRGV1- and TRGV5-expressing γδ T cells localized in the γδT17 clusters (Fig. 2D). Interestingly, V(D)J profiling of these cells revealed that TRGV1 gamma chains in the γδT17 supercluster exhibited a strong tendency for pairing with germline-encoded TRDV4 chains (Fig. S3), consistent with the identification of TRGV1/TRDV1 γδT17 in E.17 thymus (Mistri et al., 2024).

Examination of the γδT1 clusters that made up a relatively small portion of the lung γδ T cells revealed that γδT1 cells were divided into 3 distinct clusters, with C13 and C15 being composed mainly of TRGV1/TRAV15N-1 cells, while C9 was comprised mostly of TRGV4/TRDV7 γδT1 cells (Fig. 2D). All these clusters were characterized by a high expression of *Ifng, Xcl1, Ccl4,* and *Ccl5* (Fig. 2E) and the chemokine receptors *Ccr7* and *Cxcr3* (Fig. S3). The closely related C9 and C13 clusters differed in that C9 was enriched in *Eomes, Klrc1,* and *Klrc2*, while C13 was enriched in *Zfp683* (Hobit), which defines tissue-resident memory T cell precursors in the αβ T cell lineage (Parga-Vidal et al., 2021)(Suppl. Table 1). The small C15 cluster was characterized by expression of *Ifng, Gzma*, *Gzmb*, *Cd8a*, and *Cd8b1*, suggesting that it represented cytotoxic γδ T cells and that at least some of these cells might be similar to recently described CD8αβ^+^ γδT1 cells that expand in tumors (Sumaria et al., 2024) (Suppl. Table 1).

We also identified several clusters that did not segregate cleanly with either γδT1 or γδT17 transcriptional profiles. The C8 γδ T cell cluster exhibited characteristics of naïve recent thymic emigrants, and were enriched in *Cd24a, Lck, Ccr7, Sell*, and *Slamf6,* while an enrichment in cell cycle genes identified the C14 cluster as cycling γδ T cells (Fig. 2, Suppl. Table 1). The C6 cluster, which primarily consisted of TRGV6/TRDV1 and TRGV4/TRDV5-expressing cells, was enriched in ribosomal structural genes and translation factors (Fig. 2D, Suppl. Table 1), consistent with γδ T cell blasts. Finally, the C11 cluster exhibited a type I IFN response signature, while the small C12 cluster shared similarities with the C13 cluster, including expression of *Ccr8, Ccr9*, and *Cd40lg* (Fig. 2C). The C12 cluster, however, did not contain *Tbx21* or *Ifng* that are found in the C13 cluster, consistent with it being a precursor population.

Examination of the TRGV6 clusters revealed that the vast majority possessed canonical germline-encoded TRGV6 gamma chains paired with TRDV4 delta chains (i.e., Vγ6/Vδ1) that exhibited little CDR3 diversity (Fig. 2D, Fig. S2). TRGV6 clusters were characterized by a prominent C0 cluster with high expression of *Il17a, Pdcd1, Icos, Cxcr6, Il1r1, Il23r, Klrk1*, and *Areg* (Fig. 2E, Suppl. Table 1). This gene profile was consistent with a lung γδ T cell tissue-resident effector program previously described in a pre-metastatic breast cancer model (Curio et al., 2022; Edwards et al., 2023), and was consistent with the increased expression of *Areg* observed in lung γδ T cells that expand in response to Kras/p53^null^-driven lung adenocarcinoma (Jin et al., 2019). Closely related to the C0 cluster, the C2 cluster was characterized by elevated *Cd69* expression suggesting recent activation, and the C3 cluster that was enriched in *Tnfsf11* (RANKL) whose expression on γδ T cells has been demonstrated to be critical in thymic epithelial cell development (Roberts et al., 2012). The C5 cluster was characterized by enrichment of genes associated with metabolism, including *Acsbg1*, an acyl-CoA synthetase that was recently shown to modulate ST2+ Treg function (Palatella et al., 2026), as well as the sphingosine-1-phosphate receptors *S1pr1* and *S1pr4* whose expression has previously been linked to migration of Vγ6 T cells from the skin to the lungs (Papotto et al., 2026)(Suppl. Table 1).

Interestingly, we also observed that one TRGV6 γδT17 cluster, C4, exhibited hallmarks of a transitional/hybrid γδT1/γδT17 phenotype. While cells in the C4 cluster expressed the canonical *Rorc* and *Il17a* transcripts that characterize γδT17, they were also enriched in *Tbx21* and *Ifng* (Fig. 2B). In addition, in contrast to most γδT17 clusters which expressed the chemokine receptors *Cxcr6* and *Ccr6*, the C4 cluster was notably deficient in *Ccr6* expression, and was instead enriched for *Cxcr3*, a chemokine receptor that is usually associated with a γδT1 phenotype (Fig. 2C, Fig. S3). Thus, C4 γδ T cells exhibited hallmarks of recently described gut “exγδT17” cells that exhibit a transitional/hybrid γδT1/γδT17 phenotype (Baglioni et al., 2026; Parker et al., 2025).

Finally, we wanted to determine whether, in the context of a *Kras^G12D^*-expressing tumor, *Stk11*-loss differentially affected the lung γδ T cell transcriptional profile versus *Kras^G12D^* alone. We compared the hashtag-separated lung γδ T cell datasets among the KS and KRAS strains, as well as control WT and STK11 strains (Fig. 3A). This comparison revealed significant changes in cluster frequencies among the four strains (Fig. 3B). Notably, we observed a significant increase in the *Areg*-enriched C0 cluster in KS γδ T cells compared to all other strains, indicating that *Stk11* loss in lung cancer was associated with the specific expansion of *Areg*-expressing Vγ6 γδ T cells, and that *Kras* alone was not sufficient to foster this expansion (Figs. 3B-3C). Pseudobulk differential gene expression analysis revealed that, in comparison to their KRAS counterparts, KS lung γδ T cells exhibited expression of genes associated with NF-κB signaling (*Nfkbia, Nfkbiz, Nfkbid, Nfkb1*), consistent with an activated phenotype, as well as the chemokines *Cxcl10* and *Cxcl2* (Fig. 3D), the latter of which is a major chemokine involved in neutrophil recruitment (Coffelt et al., 2015; Wu et al., 2014) and is consistent with a role for γδT17 cells in recruiting neutrophils to the tumor microenvironment.

**Figure 3.**
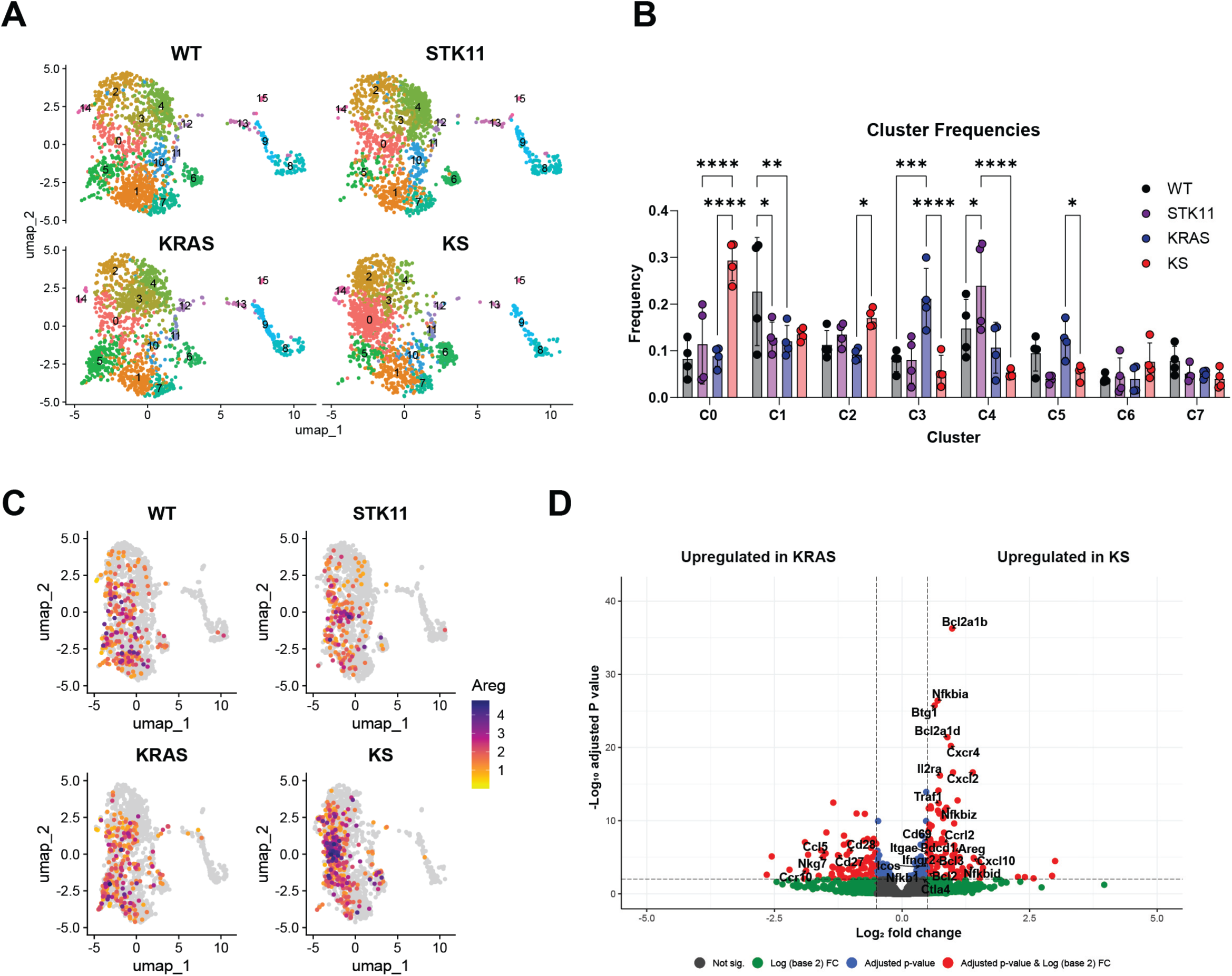
Expansion of Areg^+^ Vγ6 γδT17 in KS lungs. **A.** UMAP representation of *Kras^WT^* (WT), *Kras^WT^.Stk11^fl^* (STK11), *Kras^G12D^* (KRAS), and *Kras^G12D^.Stk11^fl^* (KS) lung γδ T cells (n = 4 each genotype), split by genotype. Numbers represent individual cluster designations. **B.** Frequencies of individual lung γδ T cell clusters from each strain as a percentage of total γδ T cells. Only the top 8 most frequent clusters are shown. **C.** Feature plots depicting γδ T cells *Areg* expression in WT, STK11, KRAS, and KS mice (purple: high, yellow: low). **D.** Volcano plot of differentially expressed genes between KS and KRAS mice; genes upregulated in KS are to the right and genes upregulated in KRAS are to the left. Genes exhibiting a log_2_FC ≥0.5 or ≤ -0.5, and a p_adj_ ≤ 0.005 are shown in red. *p < 0.05, **p < 0.01, ***p < 0.001, **** p < 0.0001.

### Expansion of lung gdT17 cells is associated with increased expression of pro-survival genes, but not increased proliferation

Next, we wanted to investigate the underlying mechanisms responsible for the expansion of the *Areg*^+^ γδT17 in KS lungs. We assessed proliferation of lung γδ T cells using an *in vivo* BrdU incorporation assay. Interestingly, we found no difference in γδ T cell BrdU incorporation between KS and KRAS lungs as well as between KS and STK11 lungs (Fig. 4A), suggesting that the significant increase in γδT17 cells is not due to increased proliferation. Consistent with this finding, an analysis of our scRNAseq data revealed no significant differences in cell cycle genes in the TRGV6^+^ cells from KS lungs compared to control mice (Fig. 4B). Moreover, we found that the *Btg1* gene, which prevents proliferation by stalling cells in the G1 phase of cell cycle (Winkler, 2010), was among the most up-regulated genes in KS lung γδ T cells (Fig. 3D). Since previous studies have shown that γδT17 cells proliferate in response to IL-1β and IL-23 (Jin et al., 2019; McKenzie et al., 2017), we assessed the levels of these cytokines in whole lung lysates and found that while KS lungs possessed significantly higher levels of IL-23 than STK11 lungs, IL-1β was nearly undetectable in either strain (Fig. 4C).

**Figure 4.**
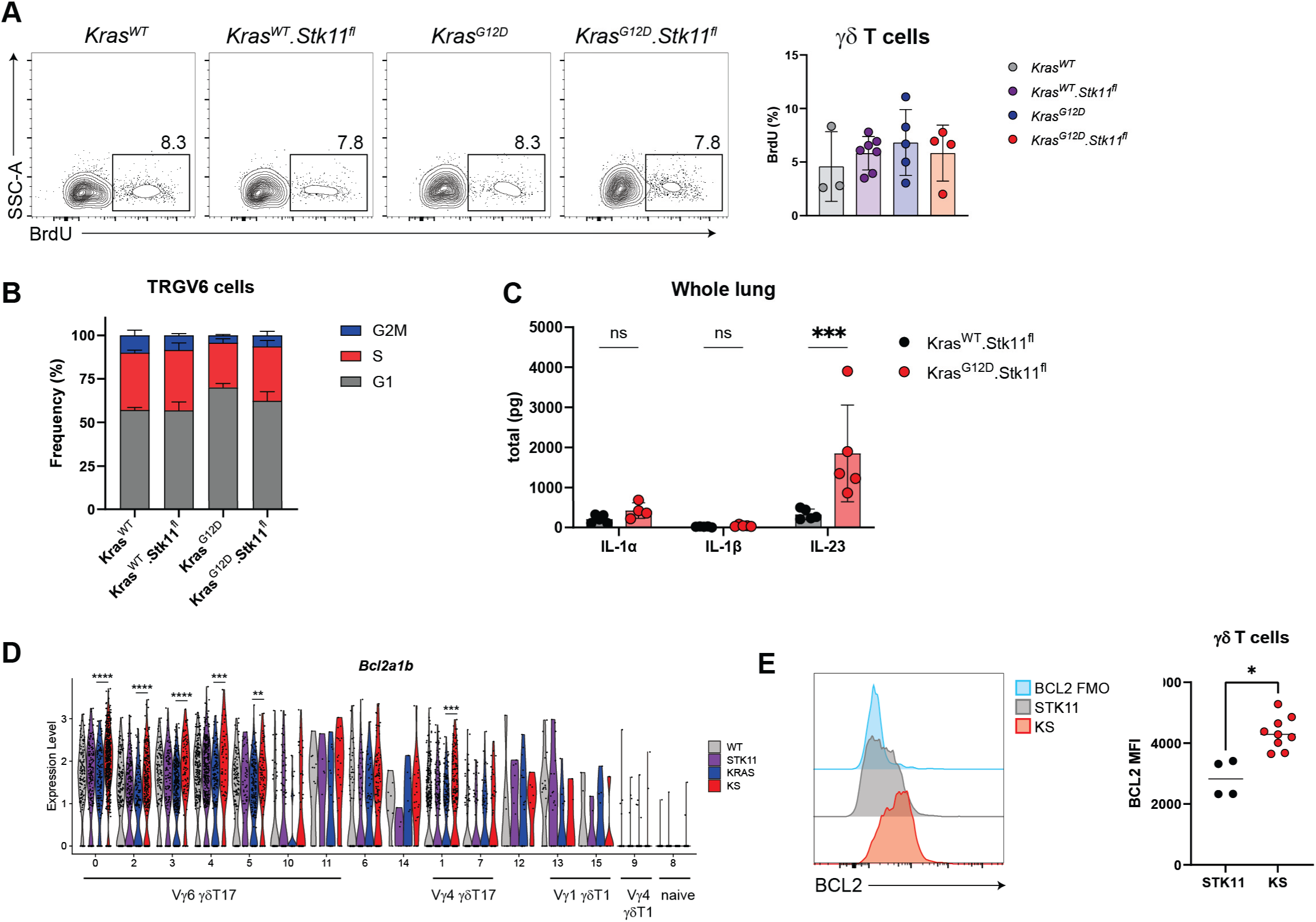
Expansion of γδT17 cells in KS lungs is associated with a pro-survival phenotype. **A.** BrdU incorporation in lung γδ T cells after tumor induction. Mice were administered a BrdU pulse on days 19 - 21 following tamoxifen induction, and BrdU incorporation was assessed using flow cytometry. Representative contour plots depicting lung γδ T cell BrdU staining *(left)* and cumulative data depicting the frequency of BrdU incorporation (*right)* are shown. **B.** Stacked bar plot depicting the frequency of lung TRGV6+ (Vγ6) cells in G1, G2M, and S phase 28 days after tamoxifen induction. **C.** Luminex analysis of IL-1α, IL-1β, and IL-23 from whole lung lysates from *Kras^WT^.Stk11^fl^* (n=5) and *Kras^G12D^.Stk11^fl^* (n=5) mice. **D.** Violin plot depicting *Bcl2a1b* expression in lung γδ clusters 28 days after tamoxifen induction in each strain; each individual dot represents a single cell. **E.** 28 days after tamoxifen induction, STK11 (n = 4) and KS (n = 9) were assessed for intracellular BCL2 expression using flow cytometry. Representative histograms (*left)* and the cumulative data depicting BCL2 MFI (*right)* are shown, *p < 0.05, **p < 0.01, ***p < 0.001, ****p < 0.0001.

Although we found no evidence of differences in proliferation between KS and KRAS γδ T cells, we did note significant enrichment in a number of Bcl2 family pro-survival genes (*Bcl2, Bcl2a1b, Bcl2a1d, Bcl2a1a,* and *Bcl3*) in KS γδ T cells (Fig. 3D, Suppl. Table 1). Examination of the expression of these genes in our single-cell dataset revealed that they were highly expressed in the TRGV6^+^ clusters C0, C2, C3, and C4, and that the expression level in all these clusters was significantly increased in KS mice (Fig. 4D). Using flow cytometry, we confirmed that KS lung γδ T cells possessed significantly higher levels of intracellular BCL2 compared to their STK11 counterparts (Fig. 4E). Thus, the mechanism underlying the expansion of *Areg*^+^*Il17a*^+^ γδ T cells in the lungs of KS mice was not due to a proliferative burst but was instead associated with the increased expression of pro-survival genes.

### *Scgb1a1*/*Kras*-driven oncogenesis results in diverse epithelial states dominated by the presence of *Krt8*+ alveolar intermediate cells

*Stk11* loss has been previously demonstrated to alter cytokine and chemokine production in alveolar type II epithelial cells (Koyama et al., 2016) in part through activation of the HIPPO/YAP1 signaling axis (Lenahan et al., 2024). Therefore, we examined whether the KS tumor cells produce cytokines or chemokines that could influence lung γδ T cell number and/or function. Since *Scgb1a1*- driven Cre is anticipated to direct tdTomato expression primarily in lung club cells in our model, we conducted scRNA-seq on FACS-sorted CD45^-^EpCAM^+^tdTomato^+^ lung epithelial cells from *Kras^WT^*, *Kras^WT^.Stk11^fl^*, *Kras^G12D^*, and *Kras^G12D^.Stk11^fl^* (n=2 each group) mice 23 days after tamoxifen induction (Fig. S4). Consistent with a previous study demonstrating that lung club cells can differentiate into other epithelial cell types during lung regeneration and tumorigenesis (Chen, 2022), UMAP clustering revealed 11 distinct clusters that could be further grouped into 6 phenotypic subsets (Fig. 5A; Suppl. Table 2). Annotation of these clusters revealed the presence of undifferentiated club cells (C3 and C9) that exhibited high levels of *Scgb1a1*, the canonical club cell marker, as well as *Scgb3a2,* a marker of variant club cells (Hewitt, 2021) (Fig. 5B-C). Mature alveolar type 2 (AT2) cells (C1, C7, C8) were also identified based on their enrichment in *Sftpc* and *Napsa*, with maturation being marked by high *Cxcl15* expression (Fig. 5B-C) (Martins, 2024). We identified an intermediate population of AT2-like progenitors (C2, C6) that retained high levels of *Scgb1a1* while expressing intermediate levels of *Napsa, Sftpc,* and *Cxcl15*, and we identified two clusters (C0 and C4) enriched in *Foxj1* and *Sntn,* consistent with ciliated epithelial cells (Fig. 5B-C) (Hewitt, 2021). Finally, we observed a subset of goblet-like cells (C10), enriched for *Muc5b, Tff1, Tff2,* and *Agr2* (Hewitt, 2021; Martins, 2024; Montoro et al., 2018), and a prominent cluster of *Krt8*^+^ alveolar intermediate cells (KACs) (C5) previously identified as a pre-malignant cell type enriched in an oncogenic *Kras* signature in both human lung adenocarcinoma and in mouse models (Han, 2024; Murray, 2022). Similar to a previous report (Han, 2024), our analysis revealed that KACs were enriched in *Cavin3*, *Plaur, Cldn4, Ndrg1*, and *Cdkn1a* (Fig. 5B-C), but lacked the *Cdkn2a* expression that characterized KACs in a *Gprc5a-/-;S*ftpcCreER model of lung adenocarcinoma (Han, 2024)(Suppl. Table 2).

**Figure 5.**
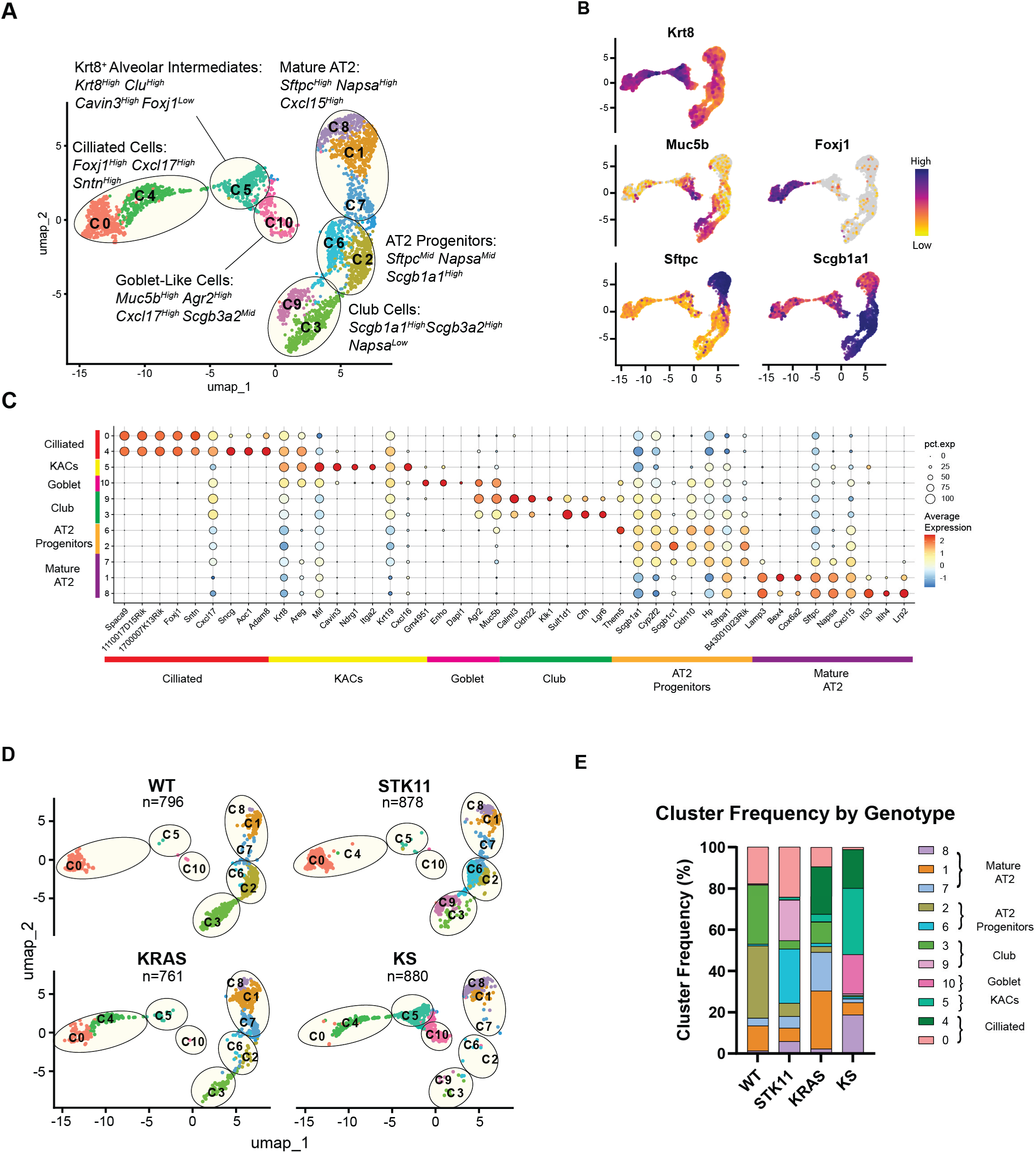
*Scgb1a1*-driven tumorigenesis yields a diverse array of lung epithelial cell subsets, including *Krt8*+ alveolar intermediate cells. **A.** UMAP visualization of FACS-sorted CD45^neg^CD326^pos^tdTomato^pos^ lung epithelial pooled from WT, STK11, KRAS, and KS lungs (n = 2 each genotype) 23 days after tamoxifen induction. **B.** Feature plots displaying relative gene expression of canonical epithelial cell type markers (high: purple, low: yellow). **C.** Dot plot illustrating the scaled expression of selected cluster-defining genes in tdTomato^+^ lung epithelium. Gene expression levels are normalized and depicted using a color scale ranging from low (blue) to high (red). The size of the dot represents the percentage of cells in the indicated cluster expressing the gene. **D.** UMAP visualization of tdTomato^pos^ lung epithelial cells, split by each genotype (n = 2). Numbers represent individual cluster designations. **E.** Abundance of distinct lung epithelial cell populations. Clusters containing the particular cell populations are colored individually. Stacked bar graph depicting frequencies of tdTomato^pos^ lung epithelial cell clusters from each strain.

To investigate the effect of the *Kras*^G12D^.*Stk11^fl/fl^*genotype versus that of the *Kras^G12D^* alone, we deconvoluted the dataset into its four genotypes and conducted pathway analysis. A comparison of differentially expressed genes between *Kras*^G12D^.*Stk11^fl/fl^*and *Kras^G12D^* cells revealed an upregulation of several pathways canonically associated with disease progression in the *Kras^G12D^*.*Stk11^fl/fl^*condition, including epithelial-to-mesenchymal transition, which in human lung adenocarcinoma is associated with both tumor progression and a more metastatic character (Schliekelman, 2015), glycolysis, Mtorc1 signaling, as well as genes associated with inflammatory response (Fig. S4). Examination of the cluster frequencies among the four genotypes revealed significant differences in the tumor subtypes associated with expression of *Kras^G12D^* alone, yielding an increase in the C1 AT2 and C4 ciliated-like clusters (Figs 5D-E). Although closely related to the C0 cluster present in the control WT and STK11 mice, the C4 cluster was enriched in *Sncg* (gamma-synuclein), *Acc1*, and *Adam8*, all of which have been reported to be overexpressed in lung cancer (Liu et al., 2005; Rocks et al., 2006; Svensson et al., 2016; Zhai et al., 2024), consistent with C4 representing a *Kras^G12D^-*driven transcriptional profile. Notably, this analysis also revealed that concurrent *Kras^G12D^* expression with *Stk11* loss-of-function induced not only an increase in the C4 cluster, but also a striking increase in the frequency of C5 KACs, C10 goblet-like cells, and C8 mature AT2-like clusters, with a corresponding decrease in the frequency of undifferentiated club, ciliated, and AT2-like progenitor cells (Fig. 5E). Taken together, these data indicate that the accelerated disease progression observed in KS mice (Fig. 1) is associated with a tumor microenvironment dominated by *Krt8*+ KACs and *Muc5b*+ goblet-like cells.

### Tumor-intrinsic *Stk11* loss leads to increased CXCL16 secretion from *Krt8*+ alveolar intermediate tumor cells

Next, we investigated whether the KACs and/or goblet lung epithelial cells that constituted KS tumors were enriched in the expression of cytokines and chemokines. This analysis revealed that several chemokines and cytokines were upregulated in KS tumors (Fig. 6A; Suppl. Table 2), including *Il6, Il33*, and *Cxcl3*, which have previously been shown to be regulated by *Stk11* expression (Koyama et al., 2016). Notably, we also observed increases in *Cxcl1* (encoding KC) that can recruit tumor-associated neutrophils to the lung via CXCR2 (Yuan et al., 2016), *Il18,* which has been reported to be present at high levels in lung cancer (Wu et al., 2022a) and has been associated with T cell exhaustion (Lutz et al., 2023), as well as *Cxcl16*, a chemokine whose levels are increased in lung cancer (Shibata et al., 2020) and has been suggested to promote proliferation and metastasis in lung cancer (Liang et al., 2018). Strikingly, in the vast majority of these cases, upregulation of the cytokines and chemokines was associated with their expression in the C5 *Krt8+* KAC cluster (Suppl. Table 2). Together, the data identify *Krt8+* KACs lung epithelial cells as the predominant source of immune-relevant cytokine and chemokine production observed in *Stk11*-null lung tumor cells in KS mice.

**Figure 6.**
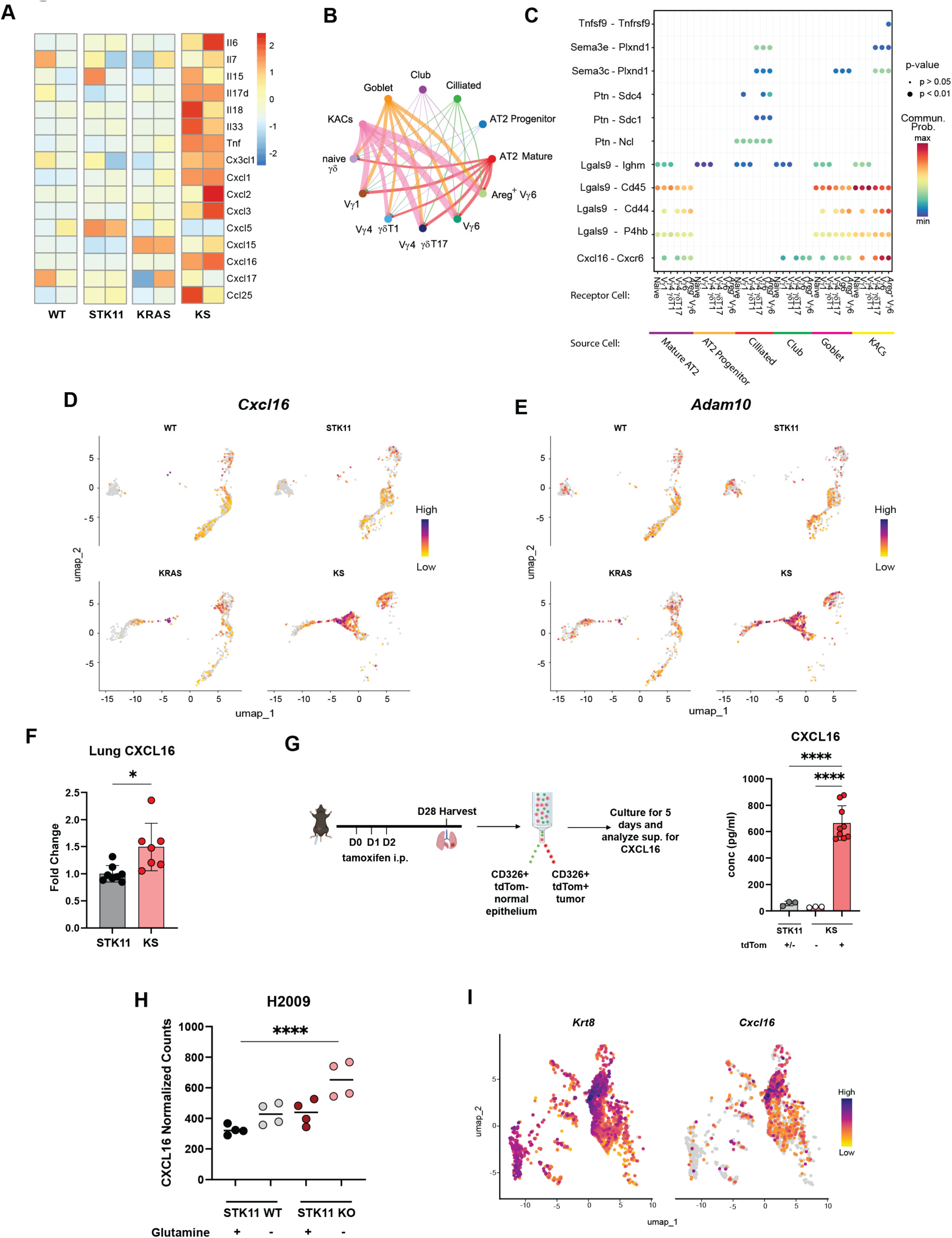
Increased CXCL16 secretion from *Krt8*+ lung alveolar intermediate cells upon *Stk11* loss. **A.** Heat map depicting selected cytokines and chemokines expressed in FACS-sorted tdTomato^+^ lung epithelial cells from WT, STK11, KRAS, and KS lungs. Results are row normalized (high: red, low: blue). **B.** Circle plot of total predicted interaction strength between lung γδ T cells and lung tdTomato^+^ epithelial cells based on gene expression of ligand/receptor pairs. The thickness of the connecting line represents the calculated strength of the interaction. **C.** Visualization of individual receptor/ligand interactions when the ligand is expressed by the lung epithelial cells and the receptor is expressed by the lung γδ T cells. The size of the dots represents the p-value, and the color of the dots represents the probability of interaction. **D.** Feature plots depicting *Cxcl16* expression in WT, STK11, KRAS, and KS lungs (purple: high, yellow: low). **E.** Feature plots depicting *Adam10* expression in WT, STK11, KRAS, and KS lungs (purple: high, yellow: low). **F.** 28 days after tamoxifen induction, STK11 (n = 8) and KS (n = 7) lungs were homogenized, and whole lung supernatant was assessed for CXCL16 using a bead-based immunoassay. **G.** 28 days after tamoxifen induction, tdTomato^+^ (n = 9) and tdTomato^-^ (n = 3) lung epithelial cells from KS mice, and total lung epithelial cells from STK11 mice (n = 3), were FACS-sorted into collagen-coated plates and expanded *in vitro* for 5 days, after which supernatants were assessed for CXCL16. Data represent the cumulative data from two independent experiments. **H.** Re-analysis of a previously published RNAseq dataset from wild-type or STK11-deficient H2009 human lung adenocarcinoma cell lines cultured with or without glutamine. Data represent normalized counts of *CXCL16* (n = 4 each group). **I.** Feature plots of previously published lung epithelial scRNAseq from a mouse model of carcinogen-induced NSCLC (Han, 2024) showing *Krt8* (left) and *Cxcl16* (right) expression. *p < 0.05, **** p < 0.0001.

We next investigated potential ligand-receptor interactions between the tumor epithelial and γδ T cells. CellChat analysis (Jin, 2024) performed on the integrated single-cell datasets predicted a high likelihood of interaction between KACs, goblet cells, and AT2 cells with γδ T cells, with KACs and goblet cells exhibiting the greatest strength of interaction (Fig. 6B). Interestingly, this analysis revealed significant interactions between KACs and Vγ6^+^ cells, especially Vγ6^+^ *Areg^+^*cells, through the *Cxcl16-Cxcr6* axis (Fig. 6C). Examination of *Cxcl16* expression in our single-cell dataset confirmed that its expression was highly enriched in the C5 KAC cluster that characterizes the KS tumor microenvironment (Fig. 6D; Suppl. Table 2).

CXCL16 can be expressed in both a soluble form and a membrane-bound complex that regulates adhesion of both T cells and MDSCs (Korbeki, 2021). While the two forms cannot be distinguished in this data set, the proteolytic cleavage resulting in the release of soluble CXCL16 has previously been shown to be mediated by the sheddase encoded by *Adam10* (Abel et al., 2004), the expression of which is also upregulated in the C5 KAC cluster (Fig. 6E). Analysis of total CXCL16 in whole-lung homogenate revealed a significant increase in total CXCL16 in KS lungs compared to that of STK11 control lungs (Fig. 6F). Together, these data suggested the possibility that the *Krt8*+ KACs that developed upon *Stk11* deletion were reprogrammed to secrete CXCL16. To test this hypothesis, we sorted total CD45^-^EpCAM^+^ epithelial cells from STK11 control lungs, as well as tdTomato^+^ (tumor cells) and tdTomato^-^ (adjacent non-tumor epithelium) CD45^-^EpCAM^+^ epithelial cells from KS lungs, cultured them *in vitro*, and assessed CXCL16 protein in the supernatant. This analysis revealed that while CXCL16 protein was nearly undetectable in the STK11 control and KS tdTomato^-^ epithelial cell supernatants, the KS tdTomato^+^ supernatants possessed significant levels of CXCL16 (Fig. 6G). Thus, KS tumor epithelial cells, but not their adjacent normal epithelial cells, secrete significant levels of the CXCL16 chemokine.

To investigate the translational relevance of CXCL16 in human lung adenocarcinoma, we reanalyzed our previously published RNAseq data of lung adenocarcinoma human cell lines in which *Stk11* was deleted using CRISPR (Lenahan et al., 2024). This analysis revealed that *STK11* deletion in human H2009 lung cancer cell lines undergoing stress induced by glutamine deprivation resulted in increased *CXCL16* expression (Fig. 6H). Next, to investigate whether increased *Cxcl16* expression in KACs could be observed in other lung cancer models, we re-analyzed a published scRNAseq dataset of epithelial cells from *Gprc5a^-/-^* mice exposed to tobacco carcinogen, which develop Kras-driven lung tumors (Han, 2024). This analysis revealed that a significant fraction of *Krt8-*expressing lung epithelial cells co-express *Cxcl16* (Fig 6I). Taken together, these analyses demonstrate that upon tumorigenesis, *Krt8*+ KACs upregulate both *Adam10* and *Cxcl16*, which together induce the secretion of significant levels of CXCL16.

### The CXCL16-CXCR6 axis promotes the expansion of lung gdT17 cells in KS mice

CXCL16 is the sole ligand for CXCR6, which has previously been demonstrated to be expressed on γδT17 (Gray et al., 2011; Wang et al., 2021b). Thus, we reasoned that increased tumor CXCL16 expression in the lungs of KS mice could be critical in driving the γδT17 expansion observed in KS lungs. Single-cell transcriptome analysis of γδ T cells confirmed that *Cxcr6* expression was highest in Vγ6^+^ cells, followed by Vγ4^+^ γδT17 cells, and that there was minimal expression in γδT1 cells (Fig. S3). We also confirmed the expression of CXCR6 via flow cytometry and found that surface protein levels mirrored the transcriptional expression patterns (Fig. 7A). To determine whether CXCL16 was playing a role in the expansion of lung γδT17 in KS mice, we conducted an *in vivo* blockade of CXCL16-CXCR6 binding. Five days after tumor induction, anti-CXCL16 was administered both i.p. and i.n. between days 5 – 20 post-induction, at which point the mice were euthanized to assess the expansion of lung γδ T cell subsets (Fig. 7B). Strikingly, blockade of CXCL16 resulted in a significant decrease in the number of lung Vγ6 cells in KS mice, suggesting that CXCL16-CXCR6 interactions were critical for the expansion of the Vγ6^+^ cells in KS lungs. Consistent with their lower level of CXCR6 expression, we also noted a downward trend in Vγ4 numbers that did not reach significance, while CXCR6^neg^ Vγ1 cells were unaffected by the CXCL16 blockade (Fig. 7C). We conclude, therefore, that CXCL16 is necessary for the expansion of lung Vγ6 cells in a model of *Kras^G12D^/Stk11*-driven lung adenocarcinoma.

**Figure 7.**
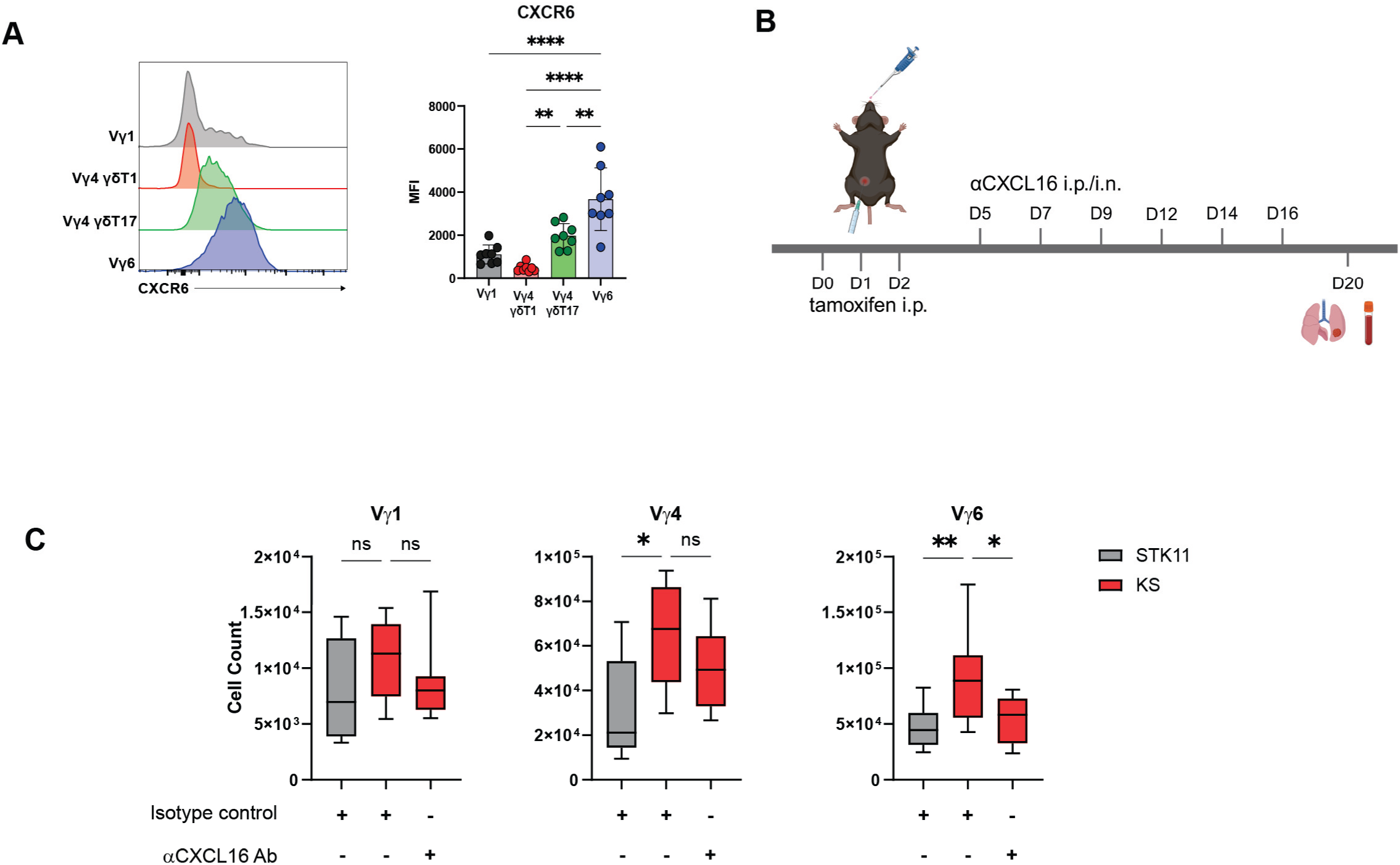
Expansion of lung γδT17 cells in KS mice is dependent on CXCL16. **A.** Representative histograms of CXCR6 expression on KS lung γδ T cell subsets *(left)* and the cumulative data representing CXCR6 MFI (*right)* are shown. **B.** Schematic of experimental design. After tamoxifen induction, mice received both i.p. and i.n. CXCL16 blockade every 2-4 days until day 20, at which time they were euthanized and lung γδ T cell counts were assessed. **C.** Cumulative data representing lung γδ T cell subset numbers from STK11 (n = 8), KS tdTomato^-^ (n = 13), and KS tdTomato^+^ mice 20 days after tamoxifen induction and administration of isotype control or CXCL16-neutralizing antibody. *p < 0.05, **p < 0.01, **** p < 0.0001.

## DISCUSSION

Although γδ T cells are now appreciated to exert significant anti-tumor and pro-tumoral effects, the factors that regulate the balance of these γδ T cell effector functions in different tumor microenvironments are still unclear. In the murine lung, the predominant pro-tumoral γδ T cells are Vγ4 and Vγ6 γδT17 cells, which drive lung tumor growth and metastasis in a *Kras^G12D^/p53^fl^* cancer model (Jin et al., 2019). In that model, lung Vγ4 and Vγ6 γδT17 cell proliferation was attributed to commensal microbiota-driven macrophage IL-1β and IL-23 production, and was consistent with the IL-1β/IL-23-mediated expansion of lung γδT17 observed in mice bearing mammary tumors (Edwards et al., 2023). Our data here provide the first scRNAseq profiling of lung γδ T cells in an autochthonous model of lung cancer and reveal that the expansion of γδT17 cells associated with increased tumor burden and rapid disease progression is associated with the specific outgrowth of an *Areg*^+^ *Il17a*^+^ subset of Vγ6 cells. In contrast to the *Kras^G12D^/p53^fl^*model, however, expansion of the *Areg*^+^*Il17a*^+^ γδ T cells did not appear to be the result of rapid proliferation but was instead the likely result of increased recruitment and/or survival cues in the lung.

Consistent with this notion, our finding of induced expression of *Bcl2a1a, Bcl2a1b*, and *Bcl2a1d* genes in the expanding *Areg+* γδT17 population is reminiscent of previous work showing that skin, but not peripheral lymph node, Vγ6 T cells express high levels of pro-survival Bcl2 family members that mediated increased protection from apoptosis, and that skin Vγ6 T cells were similarly enriched in the expression of genes encoding NF-κB proteins (Tan et al., 2019). Given that the NF-κB signaling pathway is a canonical downstream mediator of IL-1 receptor signaling and that the NF-κB pathway induces Bcl2 family member expression (Wang et al., 1999), these findings are consistent with a role for IL-1β in mediating γδ T cell expansion by promoting resistance to apoptosis. Interestingly, the failure to observe increased lung Vγ6 T cell proliferation in KS mice is in line with a recent report showing that the expansion of lung γδ T cells observed in a *Kras^G12D^/Myc^fl^* autochthonous model of lung cancer is not associated with increased proliferation (Raffo-Iraolagoitia et al., 2026). Rather, our findings indicate that the expansion of pro-tumor *Areg*^+^ *Il17a*^+^ γδ T cells in the lungs is dependent on CXCL16, a chemokine whose secretion is enhanced in tumors upon *Stk11* loss. Altogether, these findings suggest that induced recruitment and/or survival of *Areg*+ pro-tumor γδT17 in *Kras^G12D^/Stk11^null^*lung epithelial tumors contributes to the rapid disease progression observed with this tumor genotype.

Our data demonstrating that CXCL16 is required for γδT17 expansion in the lungs of KS mice is consistent with a mechanism involving increased recruitment of *Areg+ Cxcr6^+^* Vγ6 to the lung from other tissues. Indeed, CXCL16-CXCR6 interactions have previously been implicated in modulating γδT17 recruitment and function (Chen et al., 2025; Takayama et al., 2026), although only one of these studies was specific for Vγ6 T cells (Takayama et al., 2026). These reports conflict, however, with previous parabiosis experiments suggesting that Vγ6 γδT17 exhibit little tissue mobility. Limited tissue exchange was especially true for skin Vγ6 T cells (Tan et al., 2019), and while the lung γδ T cell population does apparently undergo a significant amount of exchange with other tissues, a lung tissue-resident γδT17 population enriched in *Sdc4* and *Pdcd1* was described (du Halgouet et al., 2024). These two genes are highly enriched in the C0 subset that expands in KS lungs (Suppl. Table 1), suggesting that lung *Areg^+^ Cxcr6^+^* Vγ6 cells are a tissue-resident population. Still, these studies represent Vγ6 tissue residency under homeostatic conditions, and it is not inconceivable that pro-inflammatory signals elicited in a setting such as lung cancer may alter tissue residency patterns.

An additional possibility that is not mutually exclusive from peripheral recruitment of *Areg^+^ Cxcr6^+^* Vγ6 is that CXCL16 works to position the γδT17 cells within the lung tumor microenvironment such that they receive a distinct survival signal, such as IL-7 (Li et al., 2014; Michel et al., 2012; Webster et al., 2014). This possibility is supported by a significant amount of research on the role of CXCR6 in the recruitment and/or expansion CXCR6-expressing tissue-resident memory CD8 αβ T cells. In the lung, epithelial cell- and airway macrophage-derived CXCL16 can regulate the re-positioning of CXCR6-expressing CD8^+^ T cells (Takamura et al., 2019), but CXCL16 is not essential for recruitment of these cells from the circulation (Takamura et al., 2019; Wein et al., 2019). In the setting of lung cancer specifically, CXCR6 is essential for the positioning of CD8^+^ T cells in the tumor microenvironment adjacent to dendritic cells, which deliver a pro-survival signal in the form of IL-15 (Di Pilato et al., 2021), and in the skin, CXCR6 on dermal CD8^+^ T cells is not necessary for tissue entry. Rather, CXCR6 promotes the survival and functional reprogramming to long-term tissue-resident memory cells (Heim et al., 2026). Finally, within the innate-like T cell lineages, CXCR6-deficient mice exhibit a severe reduction in liver iNKT cells that is linked to impaired survival (Geissmann et al., 2005). These published findings, together with ours here, are consistent with a model in which enhanced tumor CXCL16 promotes both the survival and recruitment of pro-tumorigenic γδT17 in the lung tumor-immune microenvironment.

Although the pro-tumoral nature of IL-17-producing γδ T cells is well described in mice, translating these findings to the clinical setting has been difficult because IL-17-producing γδ T cells are scarce in humans. IL-17-producing γδ T cells appear to be largely absent in humans under homeostatic conditions (Gray et al., 2024), and they have been described only in a few cases of chronic inflammation and infection (Hvas et al., 1993; Peng et al., 2008; Schirmer et al., 2013). However, it has recently been reported that amphiregulin-producing γδ T cells are present in human colorectal cancer, and that they can both hinder tumor infiltration and clearance by cytotoxic T cells (Harmon et al., 2023). Thus, while human tumor microenvironments may not contain abundant IL-17-producing γδ T cells, they may be enriched in pro-tumoral *Areg^+^* γδ T cells, similar to the ones described here.

A key finding of this work is the discovery that *Stk11* loss in the context of *Kras^G12D^* expression is associated with the expansion of tumor cells displaying a *Krt8^+^*alveolar intermediate cell (KAC) phenotype. *Krt8^+^* alveolar intermediate cells were first described in a mouse model of pulmonary fibrosis as an intermediate cell type in the transition from alveolar type 2 (AT2) to alveolar type 1 (AT1) cells (Strunz et al., 2020), and have been found to be associated with fibrosis (Peng et al., 2026; Strunz et al., 2020). In cancer, *KRT8* expression in tumors is associated with poor prognosis and metastasis in patients with lung adenocarcinoma (Chen et al., 2022), and KACs have been proposed as a transitionary cell state between normal epithelial cells and tumor cells and have been found to originate from both AT1 and AT2 cells (Yang et al., 2023). Strikingly, a recent report using single-cell profiling of lung epithelial cells from early-stage human lung adenocarcinoma revealed that KACs were enriched in lung precancer cells and in LUAD cells where they signified poor survival (Han, 2024). Our findings here support a role for club cells in the origin of KACs (Strunz et al., 2020), although confirmation would require additional lineage tracing experiments. Moreover, given the outsized expansion of the KACs relative to other KS lung epithelial cell types, our data suggest that loss of the *Stk11* tumor suppressor preferentially supports KAC expansion.

Our finding that CXCL16 is upregulated on KACs is consistent with previous reports demonstrating CXCL16 expression on lung cancer cells (Hald et al., 2015; Hu et al., 2014; Liang et al., 2018). The link between CXCL16 expression and prognosis in lung cancer is varied however, with some reports indicating that CXCL16 expression is associated with poor lung cancer prognosis and that it aids in tumor invasion and metastasis (Hu et al., 2014; Liang et al., 2018), while another study reported that CXCL16 expression is a positive prognostic marker (Hald et al., 2015). Mechanistically, CXCL16 was found to be essential for recruiting and positioning CTLs in various cancer types, thereby contributing to tumor control (Di Pilato et al., 2021; Matsumura et al., 2008; Muthuswamy et al., 2021; Song et al., 2025; Wang et al., 2021a). Altogether, the data reported here demonstrate that the rapid disease progression observed upon *Kras^G12D^/Stk11^null^* tumors is associated with a CXCL16-secreting *Krt8^+^* KAC tumor subtype, and that CXCL16-CXCR6 interactions are necessary to drive the expansion of pro-tumor lung *Areg^+^ CXCR6^+^* Vγ6 T cells in lung cancer.

## ACKNOWLEDGEMENTS

We thank Kris Finstaad and Kirsten Tracy in the Vermont Integrated Genome Resource for help with scRNA-seq library preparation, and Roxana del Rio Guerra and Kyra Lee for help with cell sorting. We also thank Kim Cyr and Jennie Bonica in the Office of Animal Care Management for helping maintain the mice used in these studies. This work was supported by NIH T32 AI055402 (K.J.H), NIH P30 GM118228 (J.E.B.), University of Vermont Cancer Center (J.E.B), and the University of Vermont Department of Surgery (J.E.B), NIH R21 CA280266 (D.J.S), and NIH R01 CA273238 (Y.M.W.J-H).

## Supplemental Figures

**Figure S1.**
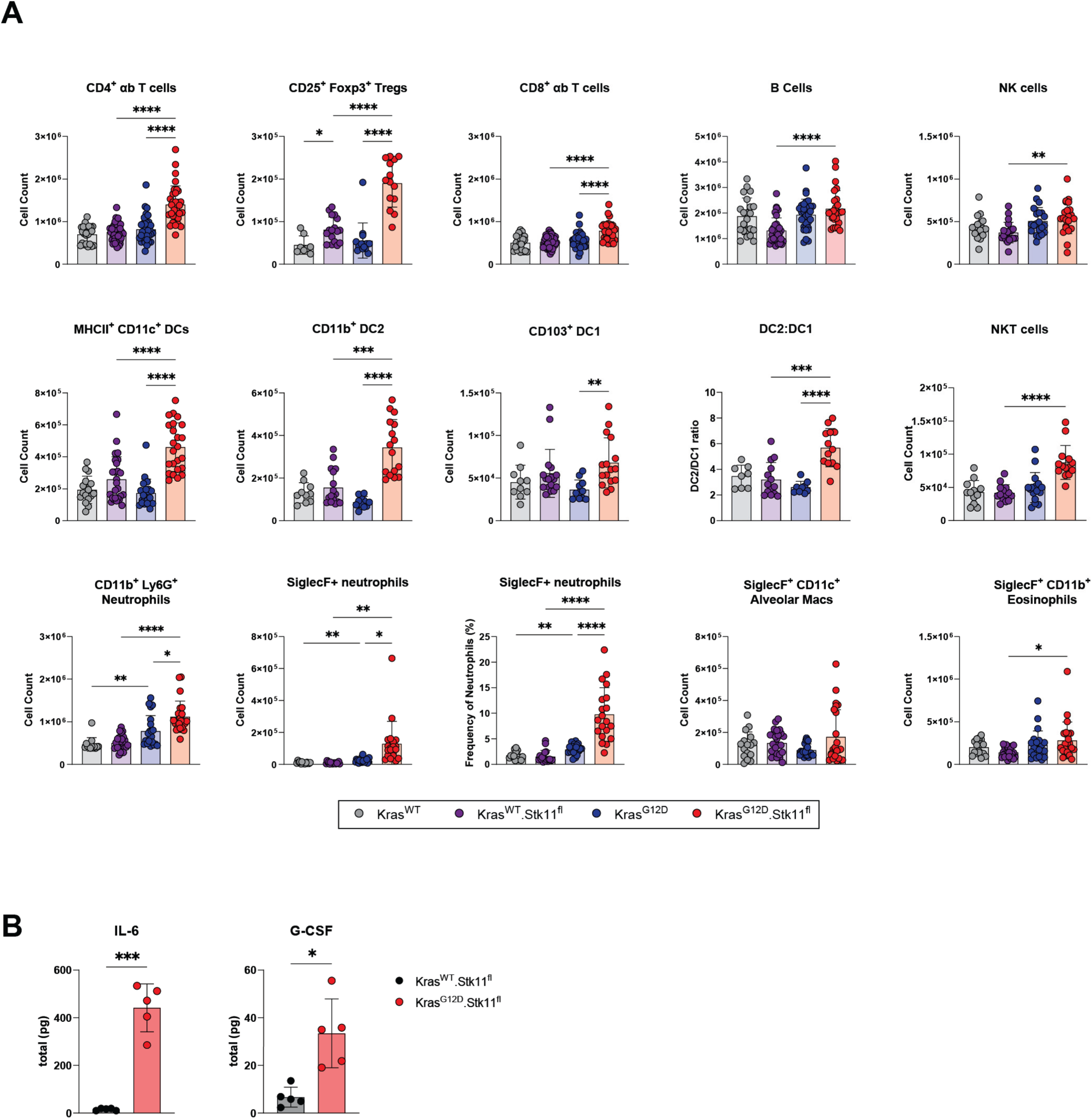
Immunophenotyping of mouse lungs after lung adenocarcinoma induction. **A.** Counts of lung leukocyte subsets in WT, STK11, KRAS, and KS lungs 28 days after tamoxifen induction, as assessed by flow cytometry. **B.** Total amounts of IL-6 and G-CSF in KS vs STK11 whole lung homogenate 28 days after tamoxifen induction. *p < 0.05, **p < 0.01, ***p < 0.001, **** p < 0.0001.

**Figure S2.**
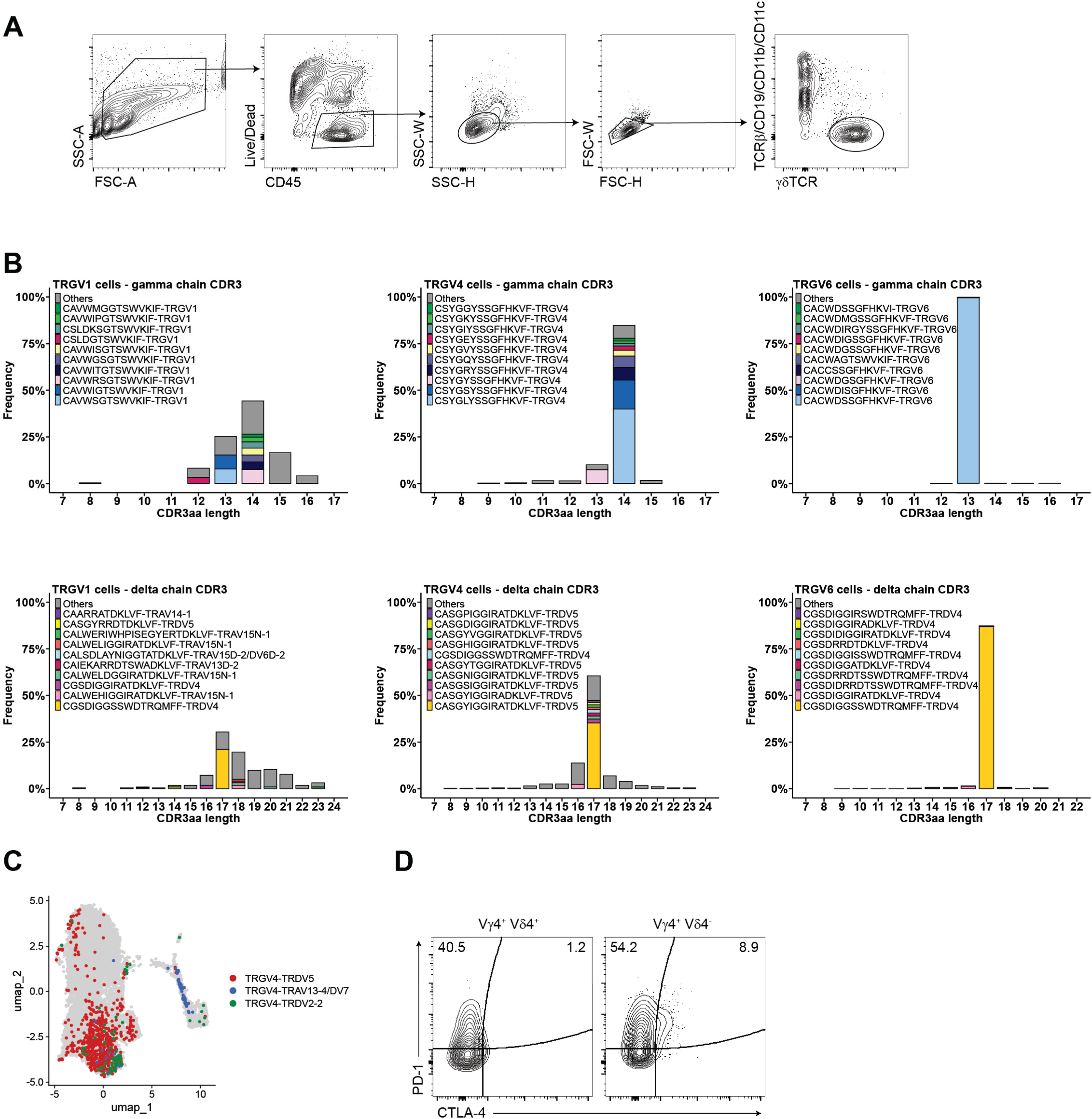
Lung. γ**δ T cell single-cell CITEseq with V(D)J profiling. A.** Representative gating scheme for FACS sorting of lung γδ T cells. **B.** Amino acid length distribution of CDR3γ (*top*) and CDR3δ (*bottom*) in TRGV1^+^ cells (*left*), TRGV4^+^ cells (*middle*), and TRGV6^+^ cells (*right)*. The top ten most frequent sequences are highlighted in color, whereas all other sequences are colored grey. **C.** UMAP displaying the distribution of different TCRγ and TCRδ pairings for TRGV4^+^ cells. **C.** Representative contour plot depicting CTLA-4 and PD-1 expression on Vγ4/Vδ4^+^ (TRDV2-2) and Vγ4/Vδ4^-^ from KS lungs.

**Figure S3.**
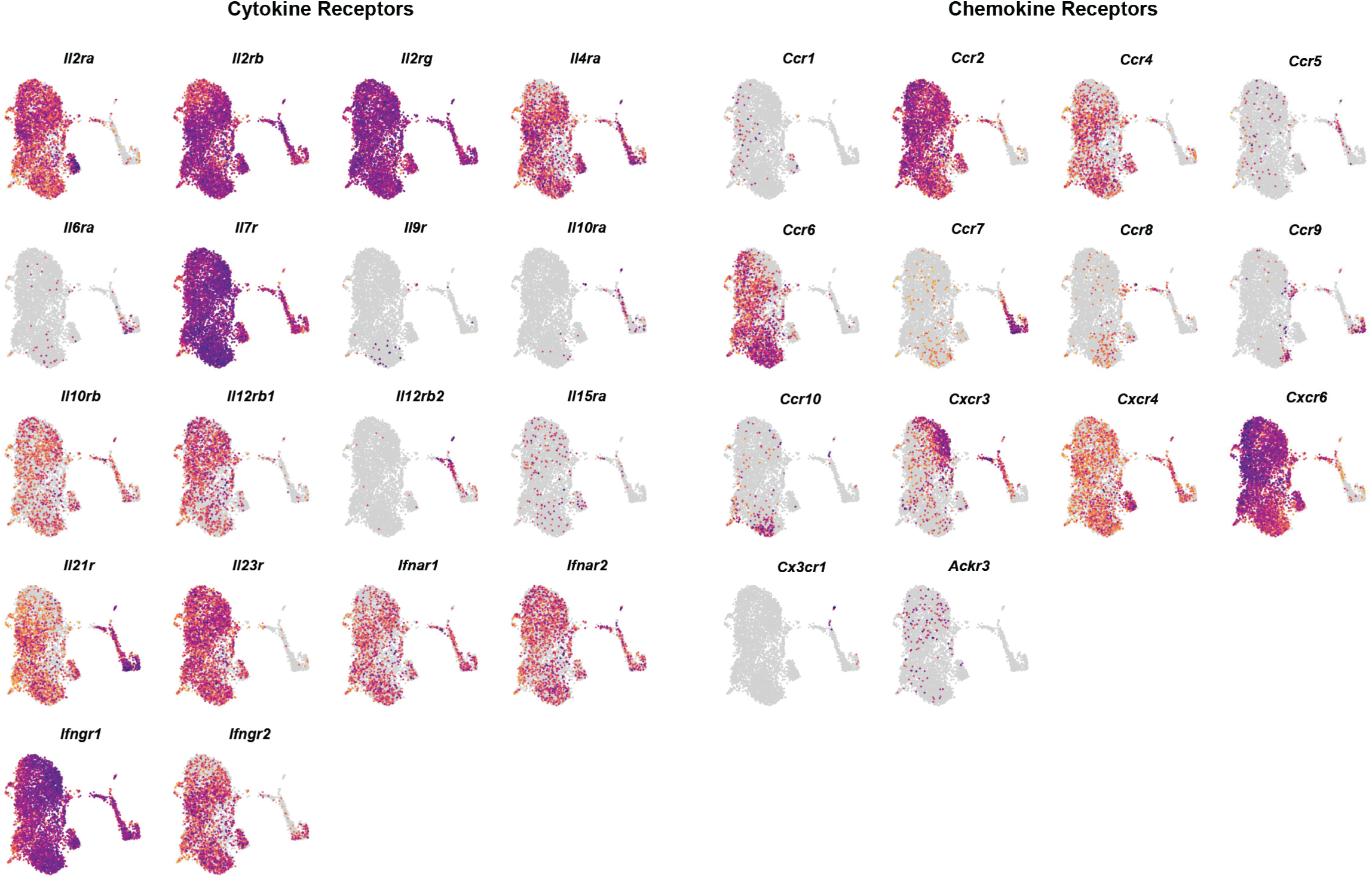
Cytokine and chemokine expression profiles of lung γδ T cells. Feature plots of lung γδ T cells from all four genotypes depicting gene expression of selected cytokine receptors (*left)* and chemokine receptors (*right*) (purple: high expression; yellow: low expression).

**Figure S4.**
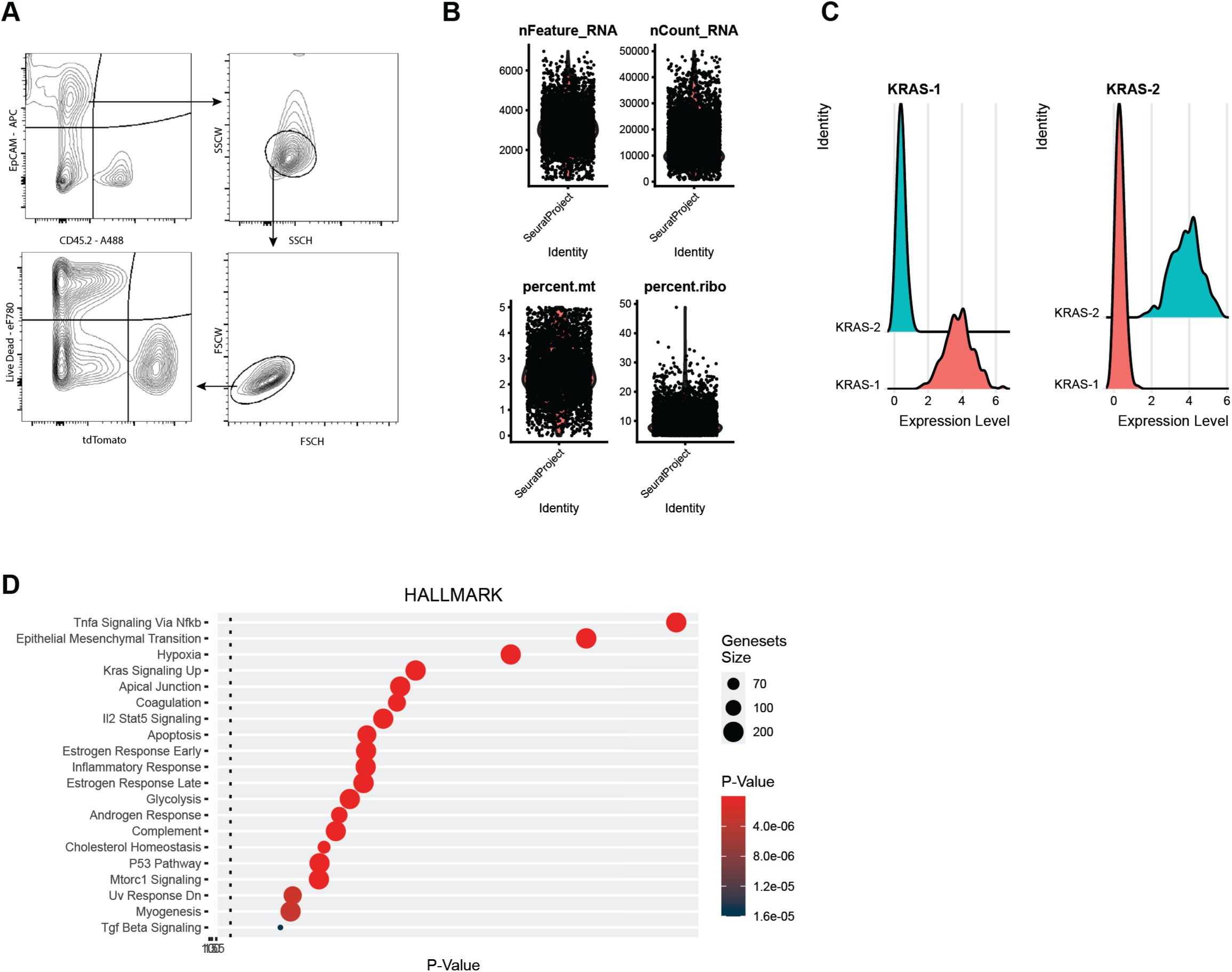
Quality control and pseudobulk pathway analysis for lung tdTomato^+^ epithelial cell single-cell RNAseq. **A.** Representative gating scheme for FACS sorting of lung tdTomato^+^ epithelial cells. **B.** Violin plots showing total unique gene (nFeature) counts per cell, total read counts (nCount) per cell, percent mitochondrial and percent ribosomal genes per cell. **C.** Ridge plots demonstrating demultiplexing for select hashtags. **D.** HALLMARK pathway analysis of lung tdTomato^+^ epithelial cells. The size of each dot represents the gene set size, and the color represents p-value.

